# Comparative genomics of bdelloid rotifers: evaluating the effects of asexuality and desiccation tolerance on genome evolution

**DOI:** 10.1101/226720

**Authors:** Reuben W. Nowell, Pedro Almeida, Christopher G. Wilson, Thomas P. Smith, Diego Fontaneto, Alastair Crisp, Gos Micklem, Alan Tunnacliffe, Chiara Boschetti, Timothy G Barraclough

**Author notes:** Current addresses Department of Genetics, Evolution and Environment, University College London, Darwin Building, Gower Street, London WC1E 6BT, UK (PA); MRC Laboratory of Molecular Biology, Francis Crick Avenue, Cambridge Biomedical Campus, Cambridge, CB2 0QH, UK (AC). Contributions These authors contributed equally to this work (Chiara Boschetti, Timothy G Barraclough).

## Abstract

Bdelloid rotifers are microscopic invertebrates that have existed for millions of years apparently without sex or meiosis. They inhabit a variety of temporary and permanent freshwater habitats globally, and many species are remarkably tolerant of desiccation. Bdelloids offer an opportunity to better understand the evolution of sex and recombination, but previous work has emphasized desiccation as the cause of several unusual genomic features in this group. Here, we evaluate the relative effects of asexuality and desiccation tolerance on genome evolution by comparing whole genome sequences for three bdelloid species: *Adineta ricciae* (desiccation tolerant), *Rotaria macrura* and *Rotaria magnacalcarata* (both desiccation intolerant) to the only published bdelloid genome to date, that of *Adineta vaga* (also desiccation tolerant). We find that tetraploidy is conserved among all four bdelloid species, but homologous divergence in obligately aquatic *Rotaria* genomes is low, well within the range observed between alleles in obligately sexual, diploid animals. In addition, we find that homologous regions in *A. ricciae* are largely collinear and do not form palindromic repeats as observed in the published *A. vaga* assembly. These findings are contrary to current understanding of the role of desiccation in shaping the bdelloid genome, and indicate that various features interpreted as genomic evidence for long-term ameiotic evolution are not general to all bdelloid species, even within the same genus. Finally, we substantiate previous findings of high levels of horizontally transferred non-metazoan genes encoded in both desiccating and non-desiccating bdelloid species, and show that this is a unique feature of bdelloids among related animal phyla. Comparisons within bdelloids and to other desiccation-tolerant animals, however, again call into question the purported role of desiccation in horizontal transfer.

## Introduction

The bdelloid rotifers are a class of microscopic invertebrates found in freshwater habitats worldwide. Two life-history characteristics make these soft-bodied filter-feeders extraordinary among animals. First, bdelloids famously lack males [1] or cytological evidence of meiosis [2, 3], and are only known to reproduce via mitotic parthenogenesis. They are therefore one of the best-substantiated examples of a eukaryotic taxon that seems to have evolved without sex or meiosis for tens of millions of years [1, 2, 4, 5]. Famously labelled as “an evolutionary scandal” [6], bdelloids have diversified into over 500 species [7, 8] in apparent defiance of the usual fate of asexual lineages [9–11]. Their persistence has implications for theories of the evolution of sex and recombination, a fundamental puzzle in biology [12–15]. A second key feature is that most bdelloid species are remarkably resilient to desiccation, and can survive the loss of almost all cellular water at any stage in their life cycle, including as adults [16, 17]. On desiccation, animals contract their body into a flat, ellipsoid “tun” shape and enter a dormant state called anhydrobiosis, during which all metabolic activities associated with life are suspended [5, 16, 18]. Individuals can remain in this condition for long periods, usually days or weeks but occasionally several years [19, 20]. The return of water restores metabolism and reproduction, with little evidence of negative fitness consequences for survivors [21]. Species that live in limnoterrestrial habitats such as puddles, leaf litter and moss are subject to cycles of drying, and the ability to survive desiccation has been proposed to play a key role in bdelloid evolution [5, 22].

Initial marker-based analyses of bdelloid genomes recovered highly divergent gene copies that were interpreted as non-recombining descendants of ancient former alleles [4]. Along with the low copy-number of vertically inherited transposable elements (TEs) [23], this result was considered positive genomic evidence of long-term asexual evolution. However, subsequent investigations of larger genomic regions revealed evidence of tetraploidy, probably arising from an ancient hybridization or genome duplication event affecting diploid ancestors prior to the diversification of the bdelloid families [5, 24, 25]. Thus, genes generally comprise up to four copies, arranged as two pairs with greater divergence between pairs (“ohnologs”, also known as homeologs in other polyploid systems) than within (“homologs”) [5, 24, 25]. Another extraordinary feature of bdelloid genomes was discovered concurrently: a remarkably high proportion of bdelloid genes show a high degree of similarity to non-metazoan orthologs, mostly from bacteria, but also fungi and plants, suggesting a rate of horizontal gene transfer (HGT) into the bdelloid genome at least an order of magnitude greater than that observed in other eukaryotes [26]. Later work confirmed that many genes originating by horizontal gene transfer from non-metazoans are expressed and functional [27].

The first whole genome sequence for a bdelloid [28] substantiated many of these findings. The degenerate tetraploid genome of *Adineta vaga* comprises homologous regions with low divergence (median 1.4% for coding sequences) and high collinearity (i.e., conserved gene order), and ohnologous regions with higher divergence (median 24.9%) but lower collinearity. The genome encodes remarkably few TEs (˜3% of the genome) but a high proportion (~8%) of HGT candidates, many of which (~20%) were found as quartets and were therefore presumably acquired prior to tetraploidization [28]. A number of unusual structural features were also reported, including a large number of breaks in collinearity between homologous regions, and the linkage of homologs on the same assembly scaffold, often arranged as genomic palindromes [28]. This assembly cannot therefore be decomposed into haploid sets, a finding that was interpreted as further evidence of long-term ameiotic evolution in *A. vaga* [28].

To what extent are the genomic characteristics of bdelloid rotifers explained by their unusual biology and ecology? An important starting point was the discovery that bdelloids can survive doses of ionizing radiation that would be lethal to nearly any other animal, owing to their ability to repair the resulting DNA double-strand breaks (DSBs) and recover from extensive genome fragmentation [29, 30]. Later experiments in *A. vaga* showed that genome fragmentation also occurs during desiccation, and led to the view that bdelloid genomes may be shaped by the need for recovering animals to repair DSBs arising from repeated desiccation [31]. It is then hypothesised that homologous gene copies are used as reciprocal templates for repairing DSBs, a process that would act to homogenise homologous regions periodically via gene conversion and select against individuals with excessive divergence, since template mismatches would disrupt DNA repair [24, 28–31]. In this scenario, the molecular consequences of desiccation are directly linked to the patterns of intragenomic molecular divergence observed in bdelloid genomes, via breakage and repair of DNA. However, the link between desiccation and DSBs is not unequivocal, and evidence from other anhydrobiotic taxa is mixed. For example, DNA integrity is largely maintained in desiccating tardigrades [32–34], but not in the chironomid insect *Polypedilum vanderplanki* [35]. In bdelloids, a key prediction is that species which undergo desiccation more frequently should experience higher rates of DSB repair, resulting in more opportunities for gene conversion and thus a lower level of homologous divergence.

A related hypothesis is that foreign DNA present in the environment may become incorporated into bdelloid genomes via non-homologous recombination during DSB repair following desiccation, resulting in a higher rate of HGT than is experienced by other eukaryotes [26, 28, 30, 31]. Evidence for high levels of non-homologous transfer inspired further suggestions that DSB-repair might similarly facilitate homologous recombination and genetic exchange between individual animals [26, 28, 30, 31]. These ideas remain controversial, however, and recent claims of evidence for DNA transfer between individuals of *A. vaga* [36] have subsequently been identified as artefacts of experimental cross-contamination [37]. A separate recent study reported a striking pattern of allele sharing among three individuals of another bdelloid species, *Macrotrachela quadricornifera,* which was interpreted as evidence of sexual reproduction via an unusual form of meiosis (similar to that of plants in the genus *Oenothera)* [38]. However, this result was not replicated in a larger study of the genus *Adineta* [39]. In the absence of clear evidence for either occasional sex or between-individual recombination in bdelloids (but without discounting either possibility), the exact nature of bdelloid recombination remains an open question.

Showing both extensive anhydrobiotic capabilities and putatively ancient asexuality, bdelloid rotifers sit at a unique junction in animal evolution. To better understand the relative contributions of these features to bdelloid genome evolution, we have taken advantage of natural variation in the capacity of species to survive desiccation, by sampling and comparing whole genomes from multiple taxa. In particular, many species in the genus *Rotaria* live in permanent water bodies and do not survive desiccation in the laboratory [17, 40]. Here, we present high-coverage, high-quality draft genomes for three species from two genera: the desiccation-tolerant species *Adineta ricciae* and the obligately aquatic, non-desiccating species *Rotaria macrura* and *Rotaria magnacalcarata*. These are compared with the published draft genome of *A. vaga.* Using a range of assembly approaches, we first confirm the conservation of degenerate tetraploidy in all species, and then test predictions regarding the effects of desiccation tolerance on intragenomic homologous divergence. We then investigate genome architecture within species, to ask whether the unusual genomic structures observed in *A. vaga* are a general feature across bdelloids. Finally, we contrast a range of genome characteristics, including homologous divergence, HGT content and repeat abundance across a wider range of animal taxa, allowing us to place some of the unique features of bdelloid genomes in a wider metazoan context.

## Results and Discussion

### Reference genome assembly and annotation

Reference genome sequences for *Adineta ricciae, Rotaria macrura* and *Rotaria magnacalcarata* were assembled using a combination of long- and short-read sequencing technologies (**S1 Table**). Kmer spectra of raw sequence reads indicated high (>100X) but variable coverage across sites in each genome (**S1 Fig**). In addition, a large proportion of low-coverage kmers indicated substantial polymorphism in the *R. magnacalcarata* raw data, most likely corresponding to population variation in the multi-individual DNA sample collected for this species (see Methods). Contaminating reads from non-target organisms were excluded by scrutinizing initial draft assemblies, and removing 3.1% of reads from the *R. macrura* dataset, 2.1% from *R. magnacalcarata,* and 6.3% from *A. ricciae* (including ~9 Mb sequences annotated as *Pseudomonas* spp.) (**S2 Fig**). Given the complex patterns of intragenomic divergence and gene copy-number observed in other bdelloid species [24, 25, 28], we adopted two assembly strategies. First, reference genomes were generated with a focus on maximum reduction of heterozygous regions and high assembly contiguity. Reference assemblies were constructed using the Platanus assembler [41] and were further reduced using Redundans [42] (**S3 Fig**, **S4 Fig**). Second, “maximum haplotype” assemblies were generated with a focus on maximum assembly of heterozygous regions. This was intended to minimise the confounding effects of assembly collapse, a phenomenon whereby homologous regions with no or low divergence are assembled as a single contig with twofold coverage relative to separately assembled regions. Further assembly metrics are provided in **S1 Data**, and all assemblies have been submitted to DDBJ/ENA/GenBank under the Project accession ID PRJEB23547^1^.

Genome metrics for *A. ricciae*, *R. macrura* and *R. magnacalcarata* reference assemblies are shown in **Table 1**, alongside those for the published assembly of *A. vaga* (accession GCA_000513175.1, hereafter referred to as the “2013” assembly) [28]. The reference assembly of *A. ricciae* spanned 174.5 Mb, comprising 4125 scaffolds with an N50 length of 276.8 kb (**Fig 1A**). The reference assemblies for *R. macrura* and *R. magnacalcarata* spanned 234.7 and 180.5 Mb over 29,255 and 20,900 scaffolds respectively, with N50 lengths of 73.2 and 53.3 kb. The proportion of undetermined bases (gaps) was low in all cases, accounting for 2.1%, 0.3% and 0.8% of the *A. ricciae, R. macrura* and *R. magnacalcarata* reference assemblies, respectively. The GC content was 35.6% for *A. ricciae*, 32.6% for *R. macrura* and 31.9% for *R. magnacalcarata;* thus %GC in *A. ricciae* is somewhat higher than in *A. vaga* (30.8%) and both *Rotaria* species. Gene-completeness was assessed by comparing sets of core eukaryotic genes to each reference assembly using CEGMA and BUSCO [43, 44]. Recovery of full-length CEGMA genes (*n* = 248) was 98%, 94% and 98% for *A. ricciae*, *R. macrura* and *R. magnacalcarata,* respectively, and gene duplication (average copy number per CEGMA gene) was 2.9,1.6 and 1.7, respectively. The equivalent recovery of a larger set of BUSCO core metazoan genes (*n* = 978) was 90% for all assemblies, with duplication scores of 2.0, 1.2 and 1.2, respectively. The equivalent completeness and duplication scores for the *A. vaga* 2013 assembly were 96% and 3 for CEGMA, and 97% and 2 for BUSCO.

**Fig 1.**
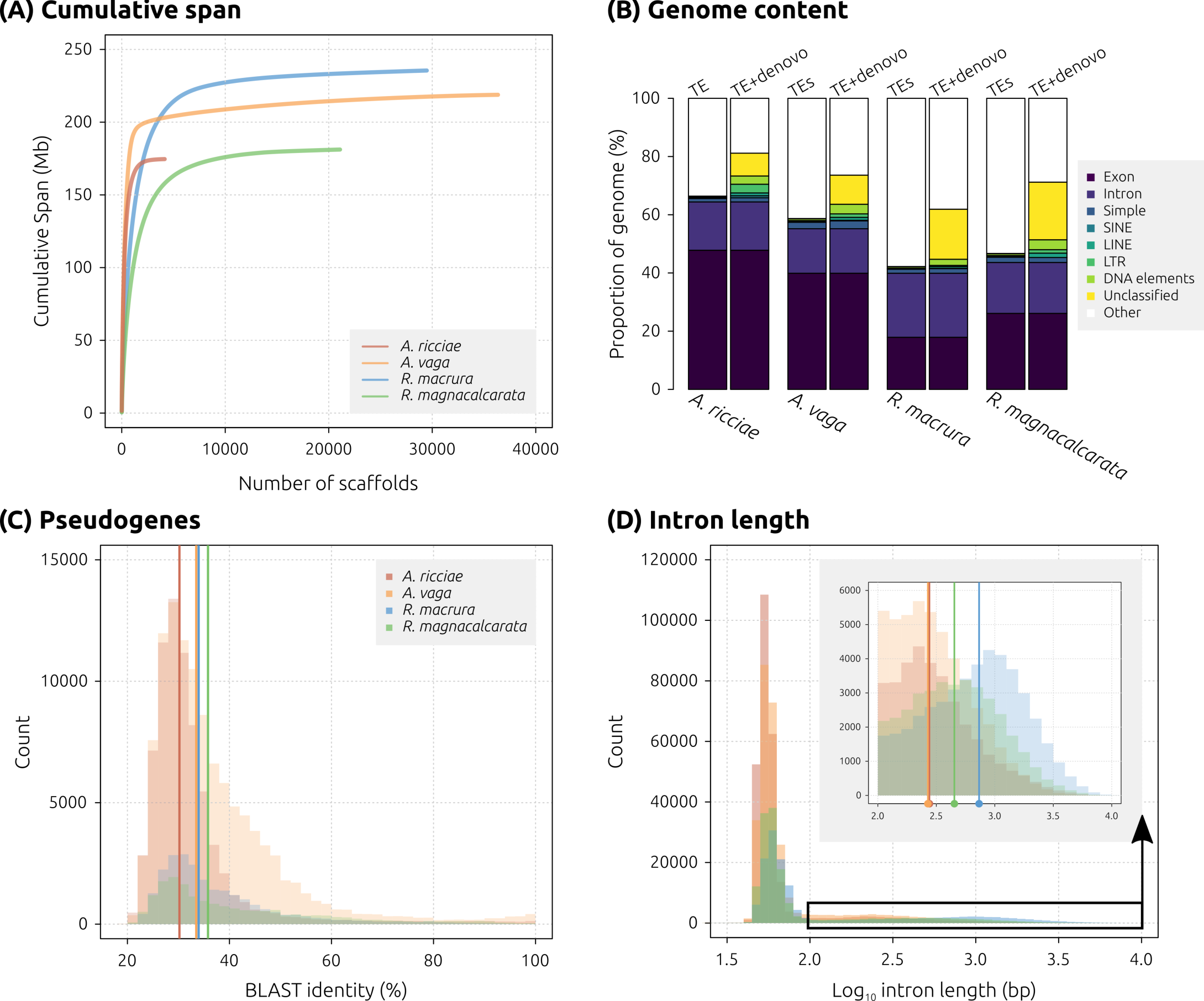
Genome properties of bdelloid rotifers. (A) Cumulative assembly span for *A. ricciae* (red), *R. macrura* (blue) and *R. magnacalcarata (green)* reference assemblies, compared to the published sequence of *A. vaga* (orange). Scaffolds are arranged in descending length order along the *X*-axis, with cumulative span plotted along the *Y*-axis. More contiguous assemblies achieve their total span with a smaller number of scaffolds, and thus manifest as a shorter line with a smaller tail. (B) Proportion of each genome covered by exons, introns, and identified repeat elements based on known metazoan transposable elements (TEs) only(left-hand column, “TE”) and TEs + de novo repeats (from RepeatModeler) (right-hand column, “TE+denovo”). Generating custom repeat libraries results in substantially greater repeat content in all species, particularly in *Rotaria.* (C) Distribution of %-identity for TBLASTN alignments (*E*-value ≤ 1e–20) of predicted proteins to their own genome, discounting hits that overlapped with any existing predicted gene model (i.e., inferred pseudogenes). Only hits with a query coverage ≥ 95% are plotted. Vertical coloured bars indicate median values. (D) Intron length distributions for each species (colours as previously). The inset shows detail of the upper tail of the main distribution (black box, truncated at ≥ 2.0). Vertical bars indicate median values for each species (of truncated distributions). Note log_10_ scale on *Y*-axis.

**Table 1.** Genome assembly and annotation metrics.

| **ASSEMBLY** |  |  |  |  |
| --- | --- | --- | --- | --- |
| **Species** | ***A. ricciae*** | ***A. vaga*<sup>1</sup>** | ***R. macrura*** | ***R. magnacalcarata*** |
| **Assembly version** | nAr.v1.8 | GCA_000513175.1 | nRc.v1.3 | nRg.v1.8 |
| **Coverage<sup>2</sup> (X)** | 134.2 | Not calculated | 210.0 | 76.3 |
| **Span (Mb)** | 174.5 | 217.9 | 234.7 | 180.5 |
| **# Contigs** | 10421 | 36335 | 31105 | 29824 |
| **Contig N50<sup>3</sup> (kb)** | 58.9 | 96.7 | 58.3 | 24.7 |
| **# Scaffolds** | 4125 | 36167 | 29255 | 20900 |
| **Scaffold N50 (kb) (#N50<sup>4</sup>)** | 276.8 (#176) | 260.3 (#240) | 73.2 (#828) | 53.3 (#890) |
| **Scaffold N90 (kb) (#N90)** | 34.5 (#868) | 7.8 (#1470) | 7.4 (#4527) | 5.3 (#4560) |
| **Scaffold longest (kb)** | 1675.3 | 1087.3 | 692.9 | 531.4 |
| **Gaps span (kb) (% genome)** | 3668.1 (2.1%) | 4136.3 (1.9%) | 640.4 (0.3%) | 1351.6 (0.8%) |
| **%GC** | 35.6 | 30.8 | 32.6 | 31.9 |
| **CEGMA**<sup>5</sup> (n = 248) | C:98%; F:99%; D:2.9 | C:96%; F:97%; D:3 | C:94%; F:98%; D:1.6 | C:98%; F:99%; D:1.7 |
| **BUSCO**<sub>EUK</sub> (n = 303) | C:97%; F:98%; D:2.1 | C:97%; F:98%; D:2.0 | C:95%; F:98%; D:1.2 | C:96%; F:97%; D:1.2 |
| **BUSCO**<sub>MET</sub> (n = 978) | C:90%; F:92%; D:2.0 | C:91%; F:92%; D:2.0 | C:90%; F:91%; D:1.2 | C:90%; F:91%; D:1.2 |
| **Reads mapped** (raw) (%) | 85.7 | Not calculated | 98.5 | 93.4 |
| **Transcripts mapped**<sup>6</sup> (%) | 99.1 | Not calculated | NA | 98.4 |
| **ANNOTATION** |  |  |  |  |
| **Species** | *A. ricciae* | *A. vaga* | *R. macrura* | *R. magnacalcarata* |
| **Method** | Braker | Maker+Augustus | Maker+Augustus | Braker |
| **# Genes** | 55801 | 67364 | 26284 | 36377 |
| **# CDS** (includes isoforms) | 58489 | 67364 | 26284 | 37380 |
| **# CDS hit to UniProt90** (Diamond E-value ≤ 1e-5) | 44974 (77%) | 49915 (74%) | 21014 (80%) | 26542 (71%) |
| **CDS mean length** (median) (bp) | 1426.0 (1059) | 1290.4 (945) | 1596.5 (1197) | 1260.7 (948) |
| **# HQ CDS**<sup>7</sup> | 49857 | 57431 | 24594 | 29359 |
| **Intron mean frequency** (introns/gene) | 4.8 | 4.1 | 5.3 | 4.0 |
| **Intron mean length** (median) (bp) | 104.1 (55) | 118.2 (58) | 362.3 (67) | 208.4 (61) |
| **Gene density**<br>(per Mb) | 319.8 | 300.3 | 112.0 | 201.5 |
| **Mean intergenic distance** (median)<br>(kb) | 1.2 (0.6) | 1.4 (0.8) | 3.9 (2.7) | 2.2 (1.4) |
| **Exon span** (Mb)<br>(%) | 83.4 (47.8%) | 86.9 (39.9%) | 42.0 (17.9%) | 47.1 (26.1%) |
| **Intron span** (Mb)<br>(%) | 29.0 (16.6%) | 33.4 (15.3%) | 51.6 (22.0%) | 31.5 (17.5%) |
| **# SEG<sup>8</sup>** (%) | 12250 (20.9%) | 16504 (24.5%) | 1149 (4.3%) | 9481 (25.3%) |
| **Transcript %GC** | 38.2 | 32.8 | 35 | 34.3 |
| **BUSCO<sub>EUK</sub>** (*n* = 303) | C:96%; F:99%; D:2.4 | C:97%; F:98%; D:2.3 | C:96%; F:98%; D:1.2 | 99%; F:100%; D:1.2 |
| **BUSCO<sub>MET</sub>** (*n* = 978) | C:92%; F:94%; D:2.4 | C:92%; F:93%; D:2.3 | C:91%; F:93%; D:1.3 | C:94%; F:96%; D:1.3 |

<sup>1</sup>Assembly metrics for *A. vaga* are based on assembly GCA\_000513175.1 by Flot *et al.* (2013) [28]. <sup>2</sup>Average read coverage, based on trimmed and filtered read data. <sup>3</sup>N50 defined as the length of the contig/scaffold at which 50% of the total genome span is reached, given a list of contigs/scaffolds in descending length order. <sup>4</sup>#N50 defined as the number of contigs/scaffolds required to reach the N50 length. <sup>5</sup>CEGMA / BUSCO notation: **C**, proportion (%) genes completely recovered; **F**, proportion (%) genes at least partially recovered (includes complete genes); **D**, duplication rate (average number of gene-copies recovered); *n*, number of queries. <sup>6</sup>Proportion of transcripts with a significant BLAT [45] alignment to the genome. <sup>7</sup>After removal of short genes with no hit to UniRef90, or hits to known transposable elements. <sup>8</sup>SEG: Single Exon Genes (genes that do not encode an intron).

The number of genes predicted from each reference assembly varied considerably among species. Gene prediction was performed using BRAKER [46] if RNASeq data were available, or MAKER + Augustus [47, 48] if not, giving initial estimates of 55,801, 26,284 and 36,377 protein-coding genes for *A. ricciae, R. macrura* and *R. magnacalcarata,* respectively (**Table 1**). Genes with BLAST matches to transposable elements (*E*-value ≤ 1e–5) were removed, and so were short genes with no matches to the UniProt90 protein database (i.e., likely spurious gene models), which resulted in “high-quality” sets of 49,857, 24,594 and 29,359 genes for downstream analyses (**Table 1**). Re-annotation of the *A. vaga* 2013 assembly resulted in 78,800 predicted genes using BRAKER or 67,364 genes using MAKER + Augustus, reducing to 57,431 after quality control. Thus, the reference genomes of *R. macrura* and *R. magnacalcarata* appear to encode approximately half the number of genes observed in *A. ricciae* and *A. vaga.* Correspondingly, the mean intergenic distance was higher in *R. macrura* (mean 3.9 kb) and *R. magnacalcarata* (2.2 kb) than in *A. vaga* (1.4 kb) or *A. ricciae* (1.2 kb) (**S5 Fig**).

We checked for mis-annotations by comparing the protein sequences to the assembly contigs from which they had been predicted, using TBLASTN (*E*-value ≤ 1e–20). This did not reveal any highly similar matches (discounting hits that overlapped with any existing predicted gene model; i.e., hits to self), indicating that “missing” *Rotaria* genes were not the result of poor gene prediction or other mis-annotation. However, the assemblies of all four species showed a large number of matches at lower similarity (30–35% median identity at the amino-acid level) (**Fig 1C**). These protein hits to non-gene annotated regions may be derived from putative pseudogenes, resulting either from degradation of coding regions following the suggested ancestral genome doubling, or more recent gene duplications that have subsequently decayed and no longer encode functional proteins.

The structure of predicted genes also varied among species. The average intron length was 104 and 108 bp for *A. ricciae* and *A. vaga* respectively, but up to three times longer in *R. macrura* and *R. magnacalcarata* (362 and 208 bp respectively) (**Fig 1D**, **Table 1**; **S6 Fig**). Distributions of intron lengths showed two distinct classes, with the majority of introns falling in the range 30–100 bp but a substantial minority showing a higher variance around a much larger mean (inset of **Fig 1D**). The mean number of introns per gene was ~4 for genes in *A. vaga* and *R. magnacalcarata*, and ~5 for *A. ricciae* and *R. macrura* (**Table 1**). The proportion of single-exon genes (SEG) was 21–25% for *A. ricciae, A. vaga* and *R. magnacalcarata,* but was substantially lower for *R. macrura* with only 4% SEG (**Table 1**).

The repeat content of bdelloid assemblies was measured using two approaches: (1) comparisons to known metazoan repeats, sampled from Repbase, (2) comparisons to Repbase plus an additional library modelled *ab initio* from each assembly, using RepeatModeler (see Methods). For (1), the relative abundances of TEs were low for all species, with the total proportion of interspersed repeats accounting for 1.2% of the assembly span for both *A. vaga* and *R. magnacalcarata,* 0.9% for *R. macrura* and 0.8% for *A. ricciae* (**S2 Data**). Including simple and low-complexity repeats resulted only in modest increases, to 2.0%, 3.4%, 2.2% and 3.0% for *A. ricciae*, *A. vaga*, *R. macrura* and *R. magnacalcarata*, respectively. For (2) however, the inclusion of *ab initio* repeats (predicted directly from the assembled nucleotides, see Methods) resulted in considerably increased repeat content for all species, but to a greater extent in *Rotaria* (16.8% for *A. ricciae,* 18.4% for *A. vaga*, 22.0% for *R. macrura* and 27.6% for *R. magnacalcarata).* A large proportion of *ab initio* repeats were marked as “unclassified” (7.8%, 10.0%, 17.2% and 19.8% of total *ab initio* repeats, respectively), and their nature is yet to be determined (**Fig 1B**; **S2 Data**). The composition of bdelloid genomes with respect to genome size evolution is considered further below.

### Marked differences in intragenomic divergence among bdelloid species

Our assembly results show an apparent twofold difference in the number of genes encoded by *Adineta* species relative to *Rotaria* species, suggesting substantial differences in either ploidy or divergence patterns between bdelloid genera. To investigate the evolutionary relationships among genes within each species, we estimated nucleotide divergence and collinearity among gene copies using MCScanX [49]. This analysis identifies collinear blocks of genes, defined as pairs of genomic regions that show conserved gene order (see Methods). We plotted the average synonymous divergence (*K*_s_) between genes within each collinear block against a “collinearity index”, defined as number of collinear genes divided by the total number of genes within a given block [28]. Both the *A.ricciae* reference and the (reannotated) *A. vaga* 2013 assemblies showed a clear delineation of genes into homologs (low *K*_s_ and high collinearity) and ohnologs (high *K*_s_ and low collinearity), as has been observed previously [28] (**Fig 2A**). Strikingly however, only ohnologous relationships are inferred in *Rotaria* genomes: collinear blocks comprised of homologous genes are not observed (**Fig 2A**). Correspondingly, the extent of collinearity was higher in *Adineta* genomes: over 75% of genes in *A. ricciae* (and 68% of genes in *A. vaga*) were categorised as members of a collinear block, compared to only 12% and 10% of genes in *R. macrura* and *R. magnacalcarata* (**S2 Table**) In addition, collinear blocks within *Adineta* species were both more numerous (1,378 and 1,828 for *A. ricciae* and *A. vaga,* respectively, versus 175 and 187 for *R. macrura* and *R. magnacalcarata*) and contained more genes (average number of genes per block was 15 and 13 for *A. ricciae* and *A. vaga*, versus 8 and 7 for *R. macrura* and *R. magnacalcarata*).

**Fig 2.**
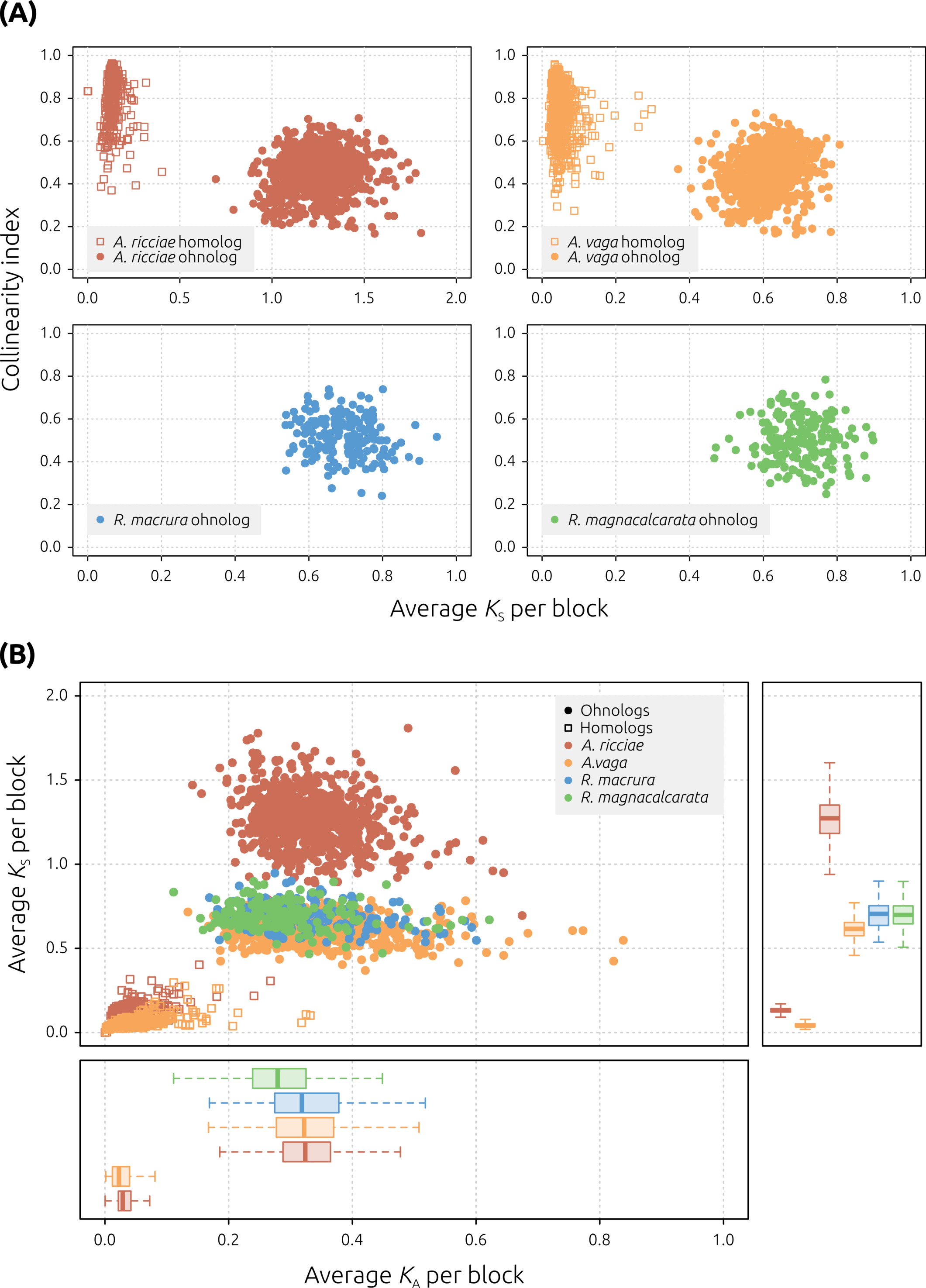
Genome collinearity within bdelloid genomes. (A) Points represent collinear blocks of genes, plotted based on the average pairwise *K*_s_ between all pairs of homologs within the block (*X*-axis) and a “collinearity index”, defined as the number of collinear genes divided by the total number of genes within the genomic boundaries of that block (*Y*-axis) (after [28]). Genes within *Adineta* species are clearly differentiated into two groups: homologs (low *K*_s_, high collinearity) and ohnologs (high *K*_s_, low collinearity), represented by open squares and filled circles, respectively. In *Rotaria* species, however, genes within identified collinear blocks only show high *K*_s_ and low collinearity scores, equivalent to that observed between ohnologs in *Adineta*. Note the different *X*-axis limit for *A. ricciae*, reflecting a higher synonymous divergence between homologs in this species. (B) Homologous genes (unfilled squares) show low numbers of synonymous (Ks) and nonsynonymous (*K*_A_) substitutions per site per block. Homologous copies are collapsed in both *Rotaria* species and are thus not shown. Ohnologous genes (filled circles) show relatively much higher rates of both *K*_s_ and *K*_A_, and are found in all species. Right-hand panel shows elevated mean *K*_s_ in both *A. ricciae* homologs (0.14 ± 0.036 (SD), versus *A. vaga* = 0.05 ± 0.026) and ohnologs (1.27 ± 0.146 versus *A. vaga* = 0.61 ± 0.062, *R. macrura* = 0.70 ± 0.080, and *R. magnacalcarata* = 0.70 ± 0.078). This elevation is not observed for *K*_A_ (lower panel). Box-plots span the median (thick line), 50% of the values (box), and 95% of the values (whiskers).

Assuming that the ancestor of extant *Rotaria* lineages was also tetraploid [25], the apparent “loss” of homologous copies in *Rotaria* species may be caused by either (1) the genuine loss of homologous gene copies from *Rotaria* genomes, resulting in a shift from tetraploidy to highly diverged diploidy, or (2) extremely low levels of divergence between *Rotaria* homologs, such that the majority of homologous sites are identical and cannot be separately assembled (i.e., are collapsed). To differentiate between these hypotheses, we characterised patterns of nucleotide polymorphism and read coverage across each genome, as has been used to investigate other polyploid genomes [50–53]. Widespread collapse of *Rotaria* assemblies is predicted to manifest as single nucleotide polymorphisms (SNPs) with a frequency around 50% and a total read-depth (reference + alternative base) that is approximately equal to the genome-wide average, analogous to collapsed heterozygous sites in a segregating diploid genome (e.g., see **Fig. 2A** of [41]). These patterns are not predicted under the hypothesis of gene loss in a separately assembled (uncollapsed) assembly, where SNPs are more likely to be the result of repetitive regions (TEs, tRNAs, low-complexity regions, etc.) or mismapped reads, and are therefore unlikely to show a frequency of 50% or consistent read-depth.

Reads were aligned to each species reference assembly, using single-clone (*A. ricciae* and *A. vaga*) or single-individual (*R. macrura* and *R. magnacalcarata*, whole-genome amplified mate-pair) libraries. The number of high-quality, biallelic SNPs that were distributed around a minor allele frequency (MAF) of 50% was 29,160 (A. *ricciae*), 27,680 (A. *vaga* 2013), 13,938 (*R. macrura*) and 52,907 (*R. magnacalcarata*), indicating at least partial collapse in all assemblies (**Fig 3A**, **S3 Data**, **S4 Data**). The relative platykurtosis observed in *Rotaria* species may be an artefact of whole-genome amplification (inflation of low-frequency SNPs), or lower coverage in general (**S7 Fig**). In *A. vaga*, the majority of sites (~76%) show coverage around 90x, representing separately assembled regions, with a minor peak at 180x, representing collapsed regions (**Fig 3B**; **S8 Fig**). The majority of SNPs occur in regions of 180x coverage, as would be expected under the scenario of localised assembly collapse [28] (**Fig 3B**). For both *R. macrura* and *R. magnacalcarata*, however, read-depth at SNP sites (reference + alternative alleles) varied in concert with the genome-wide coverage (**Fig 3B**). These patterns indicate that the majority of SNPs occur in collapsed regions, supporting the hypothesis of widespread assembly collapse in *Rotaria* species.

**Fig 3.**
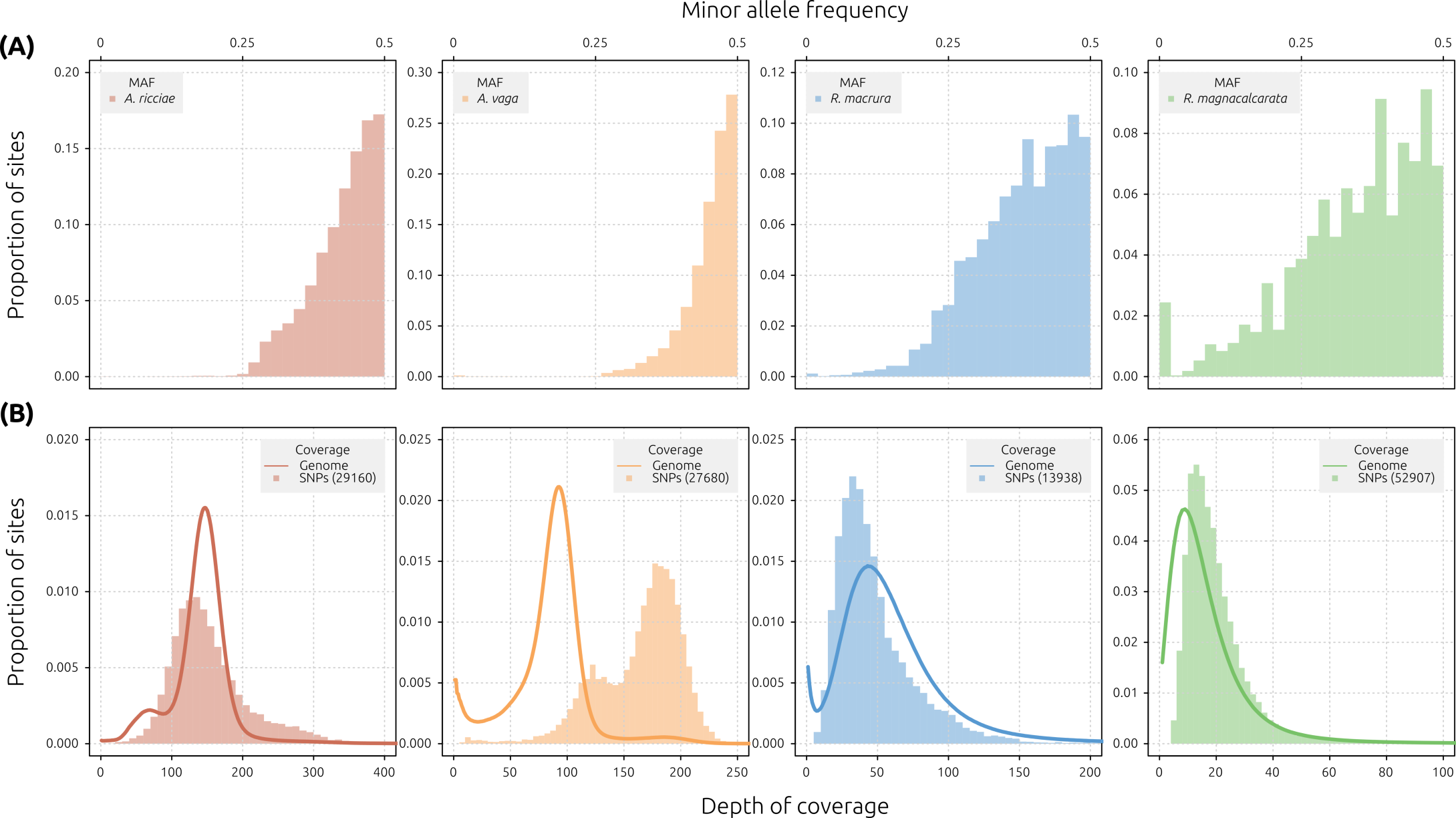
Distributions of minor allele frequency (MAF) and read coverage. Reads were mapped to reference genomes using Bowtie2 [54] and SNPs called using Platypus [55]. (A) Folded MAF spectra for detected SNPs in the reference genomes of *A. ricciae* (red), *A. vaga* (orange), *R. macrura* (blue) and *R. magnacalcarata* (green) are distributed around a mode of 0.5 in all species. (B) In each plot, the bar histogram represents the distribution of read coverage at SNP sites only, while the overlayed line shows the distribution of read coverage across all sites in the genome. The *Y*-axes indicate proportion sites with given depth of each category (i.e., peak heights are relative). The number of SNPs contributing to the bar histogram is indicated in parentheses (see legend). The cause of the secondary peak in SNP depth (at ~ 125 X) for *A. vaga* (library ERR321928) is unknown.

A different pattern is observed in *A. ricciae*, however. Here, a small proportion of sites (11%) are distributed around a peak at 75x coverage, presumably representing separately assembled regions, but the majority of sites (81%) are covered at 150x, presumably representing collapsed regions with twice the expected coverage (**Fig 3B**). Furthermore, SNP depth is unimodal and is centred around the 150x coverage peak, suggesting that most sites, including all variant sites, occur in regions of twofold coverage (150x) (**S9 Fig**). These patterns are similar to those seen in *Rotaria* assemblies, and, assuming that their interpretation is equivalent across the different species, are indicative of widespread collapse. For further context, in a diploid genome with average heterozygosity, the higher coverage peak represents collapsed regions, and SNPs occurring here are the result of successfully collapsed heterozygosity. Thus, the signal of assembly collapse that is observed in both *Rotaria* genomes is also seen in *A. ricciae*. Given that both *A. ricciae* and *A. vaga* appear to have 12 chromosomes [56, 57], this pattern is unlikely to be caused by an additional genome duplication in *A. ricciae*, but may represent an excess of tandemly duplicated gene regions (i.e., paralogs), as has been suggested to explain patterns of divergence observed for the late embryogenesis 1 gene (*LEA-1*) in *A. ricciae* [30]. Alternatively, it is possible that the *A. ricciae* sample contained cryptic population structure. Further investigations of the *A. ricciae* genome are required to test these hypotheses.

*A. ricciae* displayed a further difference from other bdelloid genomes: a clear elevation in Ks, both for homologs (compared to *A. vaga*; mean *K*_s_^Ar^ = 0.135 versus *K*_s_^Av^ = 0.05; *t* = 47, *p* < 0.01), and for ohnologs, relative to the other species (e.g. mean *K*_s_^Ar^ = 1.267 versus *K*_s_^Av^ = 0.613; *t* = 124, *p* < 0.01) (**Fig 2B**; **S3 Table**). No such elevation was observed in the rate of nonsynonymous substitution in *A. ricciae*, compared to the other species. However, the *A. ricciae* genome also shows the highest %GC of the four species (approximately 5% higher than *A. vaga*, and 3–4% higher than either *Rotaria* species). Thus, one explanation for the increase in *K*_s_ may be selection for increased G+C content in *A. ricciae*, with continued purifying selection at nonsynonymous sites [58].

### Homologous divergence in non-desiccating *Rotaria* is lower than allelic divergence in most sexuaspecies

To investigate the possibility that non-desiccating *Rotaria* species are characterised by wholesale assembly collapse, we constructed a number of uncollapsed, “maximum haplotype” assemblies (see Methods) (**S1 Data**). We also attempted to reassemble the genome of *A. vaga* for comparison, generating both collapsed and uncollapsed assembly versions. With the exception of *A. vaga*, maximum haplotype assemblies did not show the expected twofold increase in either assembly span or gene count, as would be expected if this alternative assembly approach were able to resolve homologous regions (**S2 Text, S1 Data**). Our results suggest that the choice of assembly parameters shows different overall effects in a species-specific manner, presumably depending on the extent and profile of divergence between homologous reads in each dataset (**S2 Text, S1 Data**). Overall, these results indicate that the majority of collapsed homologs in *Rotaria* species, inferred from patterns of SNP and read coverage above, cannot be separately assembled using alternative parameters. Notwithstanding confounding issues from other sources of polymorphism (e.g., the multi-individual sample for *R. magnacalcarata*), the fact that maximum haplotype assemblies for *Rotaria* species failed to recover homologous gene copies points to substantially lowered homologous divergence relative to *Adineta.*

In the *A. ricciae* reference and *A. vaga* 2013 assemblies, the majority of homologs were separately assembled, and we were able to identify homologous gene copies and estimate their sequence divergence using a BLAST-based approach [50, 53]. The median divergence between most-similar gene copies was 4.55% (mode = 3.75%) in *A. ricciae* and 1.42% (mode = 1.25%) between *A. vaga* gene copies (in close agreement with [28]) (**Fig 4A**). These estimates may be inflated however, as they fail to consider homologous regions with very low divergence that are collapsed. Alternative estimates of homologous divergence based on SNPs detected in the collapsed *A. vaga* assembly were correspondingly lower (0.955% and 0.788%, based on alignment of libraries ERR321927 and SRR801084 respectively) (**S3 Data**).

**Fig 4.**
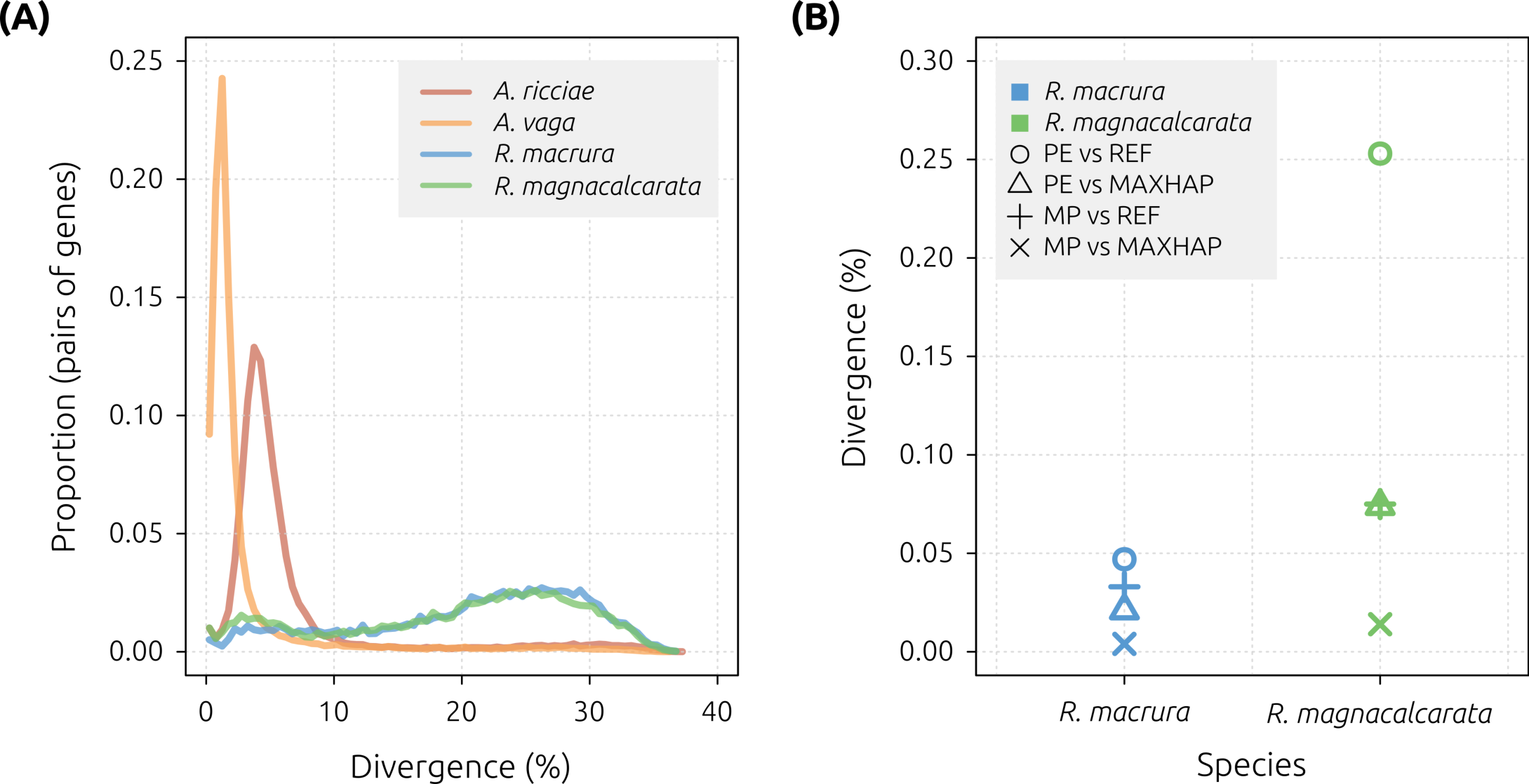
Estimates of intragenomic divergence. (A) Distribution of sequence identity (expressed as a divergence, i.e., % non-identical sites) for the top non-self BLAST hits from intragenomic comparisons within each bdelloid genome *(A. ricciae* (red), *A. vaga* (orange), *R. macrura* (blue), *R. magnacalcarata* (green)), showing highly similar gene copies within *Adineta* genomes but not in *Rotaria* genomes. Median values are 4.55% (*A. ricciae*), 1.42% (*A. vaga*), 22.5% (*R. macrura*) and 21.7% (*R. magnacalcarata*). (B) Homologous divergence estimated from SNP counts in CDS regions in reference (REF) and maximum haplotype (MAXHAP) assemblies for *R. macrura* and *R. magnacalcarata*. Mean estimates are 0.026% for *R. macrura,* and 0.104% for *R. magnacalcarata*.

To estimate the divergence between collapsed *Rotaria* homologous gene copies, we counted the number of SNPs identified previously that occurred in coding regions in *Rotaria* assemblies. Based on single-individual mate-pair libraries aligned to the *R. macrura* and *R. magnacalcarata* reference assemblies, a total of 13,115 and 36,594 SNPs were detected across 40.0 and 40.3 Mb of coding sequences (CDS), respectively (**S3 Data, S4 Data**). Assuming that all detected SNPs are the result of homologous collapse, an upper limit for the divergence between *Rotaria* homologs is thus estimated at 0.033% and 0.075% for *R. macrura* and *R. magnacalcarata* respectively (**Fig 4B**).

These results indicate that homologous divergence in non-desiccating *Rotaria* species is at least an order of magnitude lower than that observed in anhydrobiotic *Adineta* species. This contradicts hypotheses that emphasise the role of desiccation in shaping bdelloid genomes. If the rate of DSB-repair is positively correlated with the rate of gene conversion, a lower level of homologous divergence is expected in species with higher rates of desiccation. In fact, we observe the opposite: divergence between homologs in non-desiccating *Rotaria* species is considerably lower than in *A. ricciae* (median 4.6%), *A. vaga* (1.4%) (here and [28]), and *Philodina roseola* (3–5%) [24], all of which are capable of anhydrobiosis. For comparison, estimates of the sequence divergence between alleles in sexual eukaryotes range from about 0.01 % to 8% ([59]. The two *Rotaria* species fall near the lower end of this range, whereas the divergence for *A. ricciae* falls in the upper range of variation for sexual taxa.

How can we reconcile theory with these observations? One explanation is that homogenization between homologous gene copies in *Rotaria* is not caused primarily by desiccation-induced DSB-repair, but by gene conversion arising during a different process, such as mitotic crossing-over [60–62]. In the yeast *Saccharomyces cerevisiae,* for example, various forms of mitotic recombination can produce tracts of gene conversion many kilobases long, often initiated from DNA nicks that are subsequently processed into DSBs [63–65]. Such processes may be especially pronounced and irreversible in asexuals: rapid loss of heterozygosity is observed in recent invasions of asexual lineages of the water flea *Daphnia pulex*, where high rates of initial heterozygosity in hybrid asexual lineages are rapidly eroded via gene conversion and hemizygous deletion, which may ultimately limit their longevity [66].

However, this begs the question: why should the same homogenising mechanisms not operate in desiccation-tolerant species? One possibility is that desiccation-tolerant species have low or negligible background rates of gene conversion while hydrated, thanks to selection for highly effective DNA repair and error-checking mechanisms imposed by environments that desiccate on a daily basis. Such a repair system might faithfully prevent loss of diversity in the benign context of mitosis, even while occasional gene conversion remains an unavoidable consequence of the more demanding repairs required after desiccation. An alternative hypothesis is that similar homogenising forces do operate in desiccation-tolerant rotifers, but are counteracted by *de novo* point mutations generated during repair of desiccation-induced DSBs, whose net effect is to sustain high rates of homologous divergence [67]. It is not difficult to envisage other arguments, given the considerable uncertainty about the genomic consequences of desiccation. Positive evidence for a link between desiccation and DSBs is currently limited to experiments in a single species, *A. vaga* [31], and evidence from other anhydrobiotic taxa is mixed [32, 33, 35]. Further work on DNA integrity and genome evolution in bdelloids will be needed to address these divergent predictions. A final explanation that cannot be entirely excluded is that the low homologous divergence in *Rotaria* genomes results from cryptic sexual reproduction, constrained by small population sizes (i.e., inbreeding), although no males have so far been detected in *Rotaria* or any other bdelloid.

### Architectural signals of ameiotic evolution are lacking in *A. ricciae*

The *A. vaga* 2013 assembly (accession GCA_000513175.1) showed a set of unusual structural genomic features, including a large number of breaks in homologous collinearity, physical linkage of homologous genes (i.e., encoded on the same scaffold), and genomic palindromes of the form g1_A_,g2_A_,g3_A_…g3_B_,g2_B_,g1_B_, where A and B denote homologous copies of genes g1, g2 and g3. Such features would result in chromosomes that cannot be decomposed into haploid sets, and thus imply a genome architecture that is incompatible with conventional meiosis, as might be predicted given long-term asexuality [28].

To test for such structures in other bdelloid genomes, we first analysed the reannotated *A. vaga* 2013 assembly. A total of 298 breaks in collinearity (33.4% of 1,785 homologous blocks, *K*_s_ ≤ 0.3) were detected (an example is shown for scaffold AVAG00001 in **Fig 5A**). In addition, 25 homologous blocks were encoded on the same genomic scaffold, two as tandem arrays and 23 as palindromes (**Fig 5B**). Thus, our detection methods recover the same signals of ameiotic evolution reported by Flot *et al.* (2013) [28], for the same *A. vaga* assembly. The assembly method employed by Flot *et al.* (2013) was to construct contigs from Roche 454 Titanium and GS-FLX data using the MIRA assembler [68], followed by correction and scaffolding using high coverage Illumina paired-end and mate-pair data (section C1 of [28] supplement). We attempted to independently reassemble the *A. vaga* paired-end Illumina data (incorporating both mate-pair and 454 data for scaffolding) using a variety of established short-read assembly programs. However, this always resulted in highly fragmented assemblies (1–2 kb N50), except when allowing for the collapse of homologous regions (as discussed above) (**S1 Data**). The lack of contiguity in *A. vaga* maximum haplotype assemblies precluded us from using alternative assembly approaches to investigate the features detected in the 2013 assembly. A lack of separately assembled homologous gene copies in either *Rotaria* species similarly precluded structural analysis.

**Fig 5.**
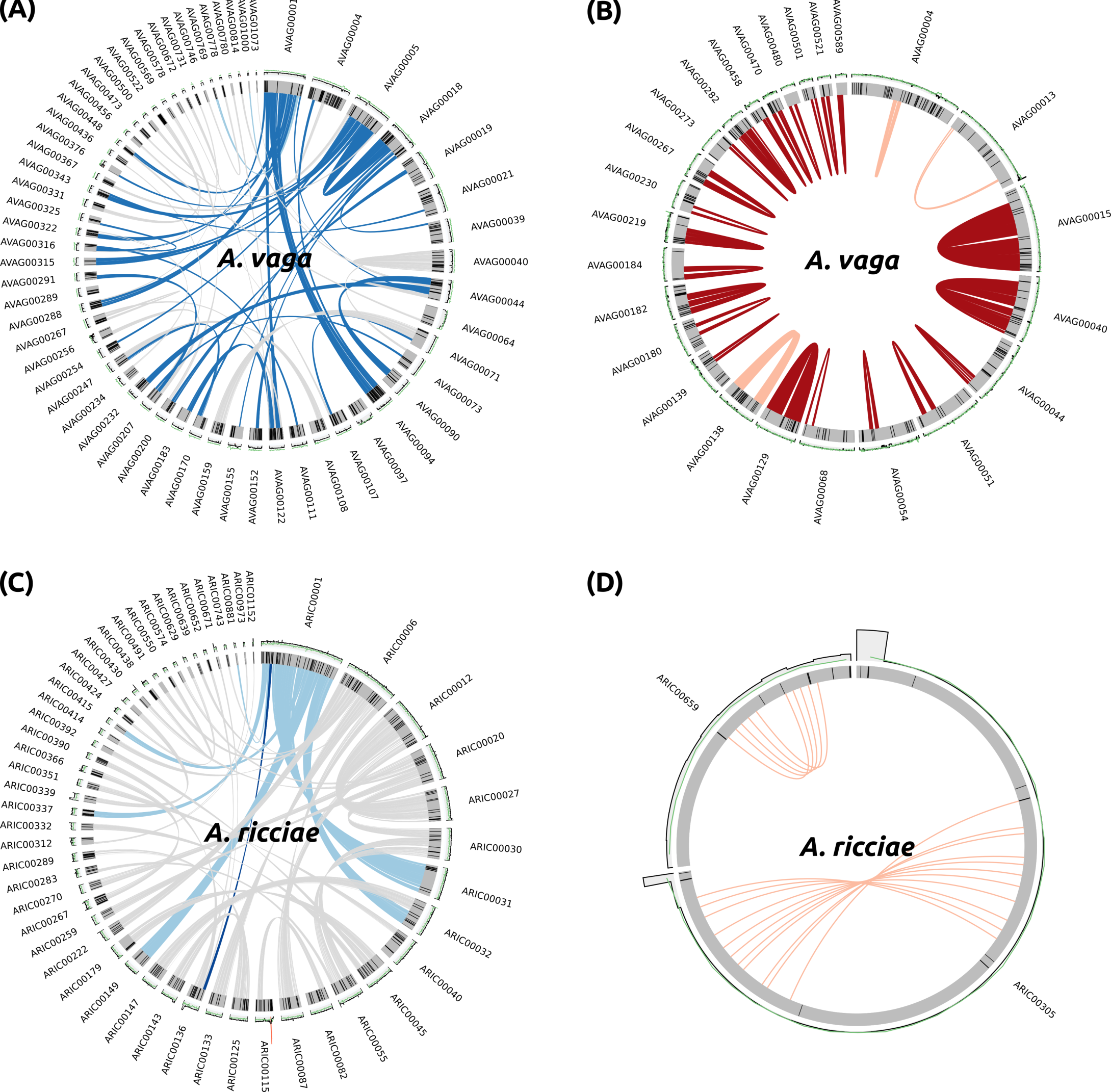
Unusual genomic features inferred from *A. vaga* are lacking in *A. ricciae.* Genomic scaffolds are shown as grey bars, %GC content (green line) and read coverage (grey histogram) averaged over 5 kb bins are shown above each scaffold. Black lines within scaffolds represent scaffold gaps (“Ns”) introduced during scaffolding of contigs. (A) Example of collinearity breaks in *A. vaga*. Homologous relationships for the longest scaffolds in both genomes (ARIC00001 and AVAG00001) are shown in light blue; downstream connections are shown in grey. Homologous blocks that represent breaks in collinearity are shown in dark blue. (B) The majority of homologous blocks encoded on the same scaffold in the *A. vaga* reference genome (GCA_000513175.1) are palindromes (dark red); tandem repeats are shown in pink. (C) Example of collinearity break in *A. ricciae.* (D) No genomic palindromes were detected in *A. ricciae*. Only two homologous blocks were detected on the same scaffold (ARIC00305 and ARIC00659), both arranged as tandem duplications.

The closely related species *A. ricciae,* however, showed both a high assembly contiguity and separately assembled gene copies. For this species, we detected only eight collinearity breaks (1.7% of 931 homologous blocks, *K*_s_ ≤ 0.5) between homologs (**Fig 5C**), in contrast with hundreds inferred from the *A. vaga* 2013 assembly. Assuming that the *A. ricciae* and *A. vaga* 2013 assemblies are accurate representations of the genome of each species, collinearity appears to be markedly more conserved between homologs in *A. ricciae* than in *A. vaga*. However, many of the detected breaks in both species span regions separated by numerous scaffold gaps (“Ns” introduced during the joining of contigs). This suggests that at least some detected collinearity breaks may be the result of errors introduced during the scaffolding process, despite requisite care taken during assembly scaffolding (**Fig 5A,C**). For example, an unscaffolded *A. ricciae* assembly showed only a single break in collinearity, although the increased fragmentation of this assembly (N50 = 18.7 kb) may limit our ability to detect such breaks in this case. In addition, we did not detect any cases of homologous genes arranged as palindromes in *A. ricciae*. Only two cases of linked homologous blocks of genes were detected for *A. ricciae*, and both were arranged as tandem repeats rather than palindromes, on relatively short scaffolds involving a small number of genes (**Fig 5D**).

Overall, these results suggest that certain unusual genomic features, previously interpreted as positive signatures of long-term ameiotic evolution in *A. vaga* [28], are largely absent from the closely related *A. ricciae* (and remain untested in other bdelloids). These patterns may reflect true differences between *Adineta* species, although no marked dissimilarity in karyotype is evident [56, 57]. Alternatively, they may be either false-positive or false-negative artefacts of applying alternative assembly methodologies to complex genomes with different patterns of intragenomic divergence.

Evidence from other taxa is limited. For example, similar features have been reported in the recently assembled genomes of the parthenogenetic springtail *Folsomia candida* [69] and in certain apomictic species of *Meloidogyne* root-knot nematodes [70]. However, these involved only small proportions of the respective genomes, and the latter study used the same assembly approach that was applied to *A. vaga* [70]. It may be noted that similar structural variants are readily detected in many sexual organisms too, including humans [71, 72], cichlid fishes [73] and cows [74]. These often involve translocations or duplications of sequences many kilobases in length. Thus it is difficult to ascertain the evolutionary significance of these features, particularly when only small proportions of the genome are involved. Even if real, they may reflect an unremarkable background level of genomic rearrangement that is independent of reproductive mode. Further work is required to improve and validate assembly contiguity using long-read technologies and physical mapping techniques.

### No evidence for widespread loss of sex-related genes in bdelloids

To investigate other potential genomic signals of asexuality, we tested for the presence of a suite of 41 sex-related genes in each bdelloid reference genome using both TBLASTN (comparing to the genome) and HMMER (comparing to the proteome). Genes were taken from a recent study by Tekle *et al.* (2017) [75], and included 11 meiosis-specific genes (*SPO11*, *REC8*, *DMC1*, *MND1*, *HOP2*, *HOP1*, *MSH5*, *MER3*, *MSH4*, *ZIP1* and *RED1*), 19 genes involved in recombinational repair, six genes involved in the sensing of DNA damage, four genes involved in DSB-repair via nonhomologous end-joining and one gene involved in bouquet formation [75] (**S6 Data**). For comparison, we also tested the genomes and proteomes of two additional species, *D. melanogaster* and *C. elegans*. Overall, a positive match using TBLASTN and/or HMMER was recorded in at least one bdelloid species for all tested genes (40 of 41, 98%) with the exception of *RED1*, involved in crossover regulation and not detected in any bdelloid at any significance threshold (**S6 Data**). However, *RED1* was not detected in *D. melanogaster* and only as a poor match in *C. elegans*, and thus may represent an ancestral loss that predates the bdelloids. Two other genes showed variable patterns of presence and/or matches at lower thresholds: *BRCA2*, involved in recombinational repair and detected only in *R. magnacalcarata*, and *HOP2*, involved in crossover regulation and detected only below the inclusion threshold in either *Adineta* species (**S6 Data**). However both genes were detected at lower thresholds in other species, and may therefore represent matches to divergent *BRCA2* and *HOP2* genes in bdelloid genomes.

Using a similar approach, we also tested for the presence of the *Zona pellucida-like* domain (Pfam accession PF00100), involved in the binding of sperm on the surface of egg cells in other animals [76] and reported as missing from the *A. vaga* genome [28]. Alignment of the domain to the *A. vaga* protein set (using HMMER) returned no hits, confirming the previous report, but alignment to the *A. ricciae*, *R. macrura* and *R. magnacalcarata* proteomes returned positive matches in all cases.

Overall, these findings suggest that bdelloids encode the majority of genes involved in meiosis and sex-related functions, including domains associated with sperm-binding. However, the presence of these genes does not necessarily indicate sex or meiosis, as genes are likely to be retained for other functions related to homologous recombination and DSB repair [77].

### Non-desiccating *Rotaria* species have larger genomes

Assuming that *Rotaria* assemblies do indeed reflect collapsed homologs, estimations of other global genome properties can be refined. The maximum haplotype assembly for *A. ricciae* was approximately 201 Mb in length encoding 63,000 genes, while reannotation of the partially collapsed 217 Mb *A. vaga* 2013 assembly showed 60,500 genes. While maximum haplotype assemblies for *A. vaga* were highly fragmented (and therefore poorly annotated), a collapsed *A. vaga* assembly, reduced to 109 Mb, encoded 31,600 genes. These values suggest that the full complement of genes (equivalent to ~4n, given degenerate tetraploidy) in *A. ricciae* may be in the region of 60,000–65,000 genes, across a total span of at least 200 Mb. The genome of *A. vaga* is probably larger, at least 240 Mb, encoding ~68,000 genes. The largely collapsed reference assemblies of *R. macrura* and *R. magnacalcarata* showed approximately 25,000 and 35,000 genes respectively, and thus are in broad agreement with observations from *Adineta*. This also implies that the total genome size for *Rotaria* is in the region of 400–500 Mb. Further evidence of genome size variation among bdelloids is supplied by comparisons to the collapsed *A. vaga* assembly, which is of equivalent ploidy but reduced by 60 Mb compared to *R. magnacalcarata* and 113 Mb compared to *R. macrura*.

What explains the observed differences in genome size? Based on comparisons to known metazoan TEs from Repbase, the abundance of TEs and low-complexity repeats was low in all genomes (2.0%, 3.5%, 2.3%, and 3.1% for *A. ricciae*, *A. vaga* 2013, *R. macrura* and *R. magnacalcarata*, respectively; **Fig 1B**, **S2 Data**), suggesting that expansions of known TEs or simple repeats in the *Rotaria* lineage is unlikely to be a major driver. However, the inclusion of *ab initio* repeats, modelled from the assembly sequences directly using RepeatModeler (see Methods), resulted in a marked increase in total repetitive sequences for all species (16.8%, 18.4%, 22.0%, and 27.6%, respectively; **Fig 1B**, **S2 Data**). The majority of *ab initio* repeats were annotated as “unclassified” and thus are not similar to known repetitive elements (in Repbase) (**S2 Data**). The relative increase is greatest in the two *Rotaria* species, suggesting that a substantial fraction of the *R. macrura* and *R. magnacalcarata* reference assemblies (~17% and 20% respectively) are covered by repeats whose exact nature remains to be elucidated. In addition, average intron sizes in *Rotaria* genes are longer (by at least 100%), driven primarily by an increase in the number of long introns (i.e., ≥ 100 bp). A similar expansion of intron size has been recently observed in genome comparisons between the tardigrade species *Hypsibius dujardini*, with a genome size of 104 Mb, and *Ramazzottius varieornatus*, with a much smaller genome size of 56 Mb [34, 78]. Intriguingly, *R. varieornatus* is capable of rapid anhydrobiosis while *H. duajardini* survives desiccation only under certain conditions, suggesting a possible link between desiccation tolerance and genome compaction in both bdelloids and tardigrades.

### High HGT content is a unique feature of bdelloid genomes

To gain a better understanding of how bdelloid genomes compare in a wider metazoan context, we characterised an additional 13 species from across the Protostomia (**S4 Table**) based on genome size, gene density, patterns of orthologous gene clustering, HGT content and repetitive sequence content (**Fig 6**, **S10 Fig**, **S5 Table**; **S6 Table**). Our comparison included a broad taxonomic range of species from different ecological niches, including a number of molluscs [79–82], annelids [80], the platyhelminth *Schistosoma haematobium [83],* the desiccation-tolerant tardigrade *R. varieornatus* [34], the reduced genome of the orthonectid intracellular parasite *Intoshia linei* [84], and chromosomal-level reference genomes for *Caenorhabditis elegans* [85] and *Drosophila melanogaster* [86]. Phylogenetic relationships among species were not estimated directly but inferred from the literature [84, 87, 88].

**Fig 6.**
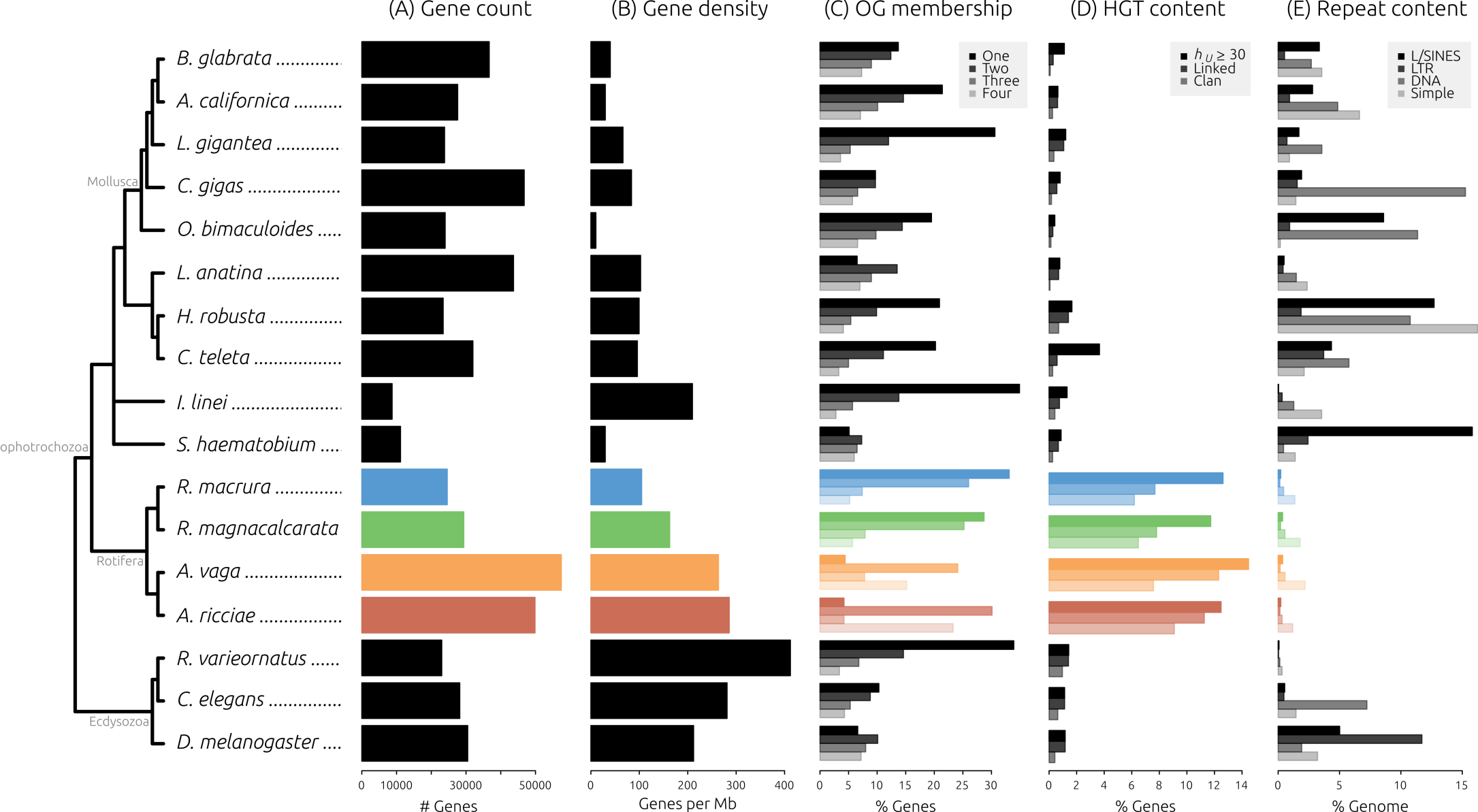
Bdelloid genome characteristics compared to other metazoans. Phylogenetic relationships among 17 protostome species are shown on the left (inferred from [84, 87, 88]), major groups are marked in grey. (A) Gene count, defined as the total number of CDS encoded in each genome. (B) Gene density, defined as the number of CDS divided by the genome (haplome) span. (C) Same-species OG cluster membership, with bars from top to bottom representing 1, 2, 3 and 4-member clusters. (D) HGT content, with bars from top to bottom representing the % of each species genes showing *h*_U_ ≥ 30 (HGT candidates, HGT_C_), “Linked”, HGTc in physical linkage with a known metazoan gene, and “Clan”, HGTC linked to metazoan genes and with clan membership to non-metazoan genes from phylogenetic analyses. (E) Repeat content, with bars from top to bottom representing the % genome covered by LINES+SINES, LTR elements, DNA elements, and simple/low-complexity repeats.

We assessed the extent of horizontal transfer into protostome genomes using both sequence comparison and phylogenetic approaches. The extent to which HGT contributes to the genomes of multicellular eukaryotes is controversial. For example, a recent claim of 17% non-metazoan genes encoded in the genome of the tardigrade *H. dujardini* was later shown to be derived mostly from contaminating non-target organisms [89–92]. Nonetheless, a high proportion of genes from a variety of non-metazoan sources has been consistently reported in bdelloid genomes from a range of independent data, including fosmid sequences [26], transcriptomes [27, 40], and whole genome data [28]. To measure the level of horizontal transfer into bdelloid genomes, we developed a HGT assessment pipeline that uses both sequence comparison and phylogenetic approaches to build a body of evidence for the foreignness of each predicted gene. Our goal was not to unequivocally assert the evolutionary history of individual genes, but rather to apply these tests consistently across the set of protostome genomes as a fair comparison for estimating HGT.

Our initial screen identified 6,221 (12.5%), 8,312 (14.5%), 3,104 (12.6%) and 3,443 (11.7%) genes from *A. ricciae*, *A. vaga*, *R. macrura* and *R. magnacalcarata* respectively as HGT candidates (denoted “HGT_C_”) (**Fig 6D**, **S5 Data**). These values are substantially higher than the proportion of HGT_C_ observed in any other protostome species included in this analysis, using the same pipeline and thresholds (the highest proportion of HGTc for a non-bdelloid was 3.6%, for the annelid worm *Capitella teleta*). This is also noticeably higher than estimates based solely on the Alien Index (**S5 Data**). For each HGT_C_ we then assessed (a) the presence of predicted introns, (b) scaffold linkage to another gene of unambiguous metazoan origin, (c) presence on a scaffold that encodes a high HGT_C_ proportion, (d) membership within a “clan” of non-metazoan orthologs, and (e) monophyly of the HGTc with all present non-metazoan orthologs to the exclusion of all metazoan orthologs (see Methods) (**S5 Data**). Testing for clan membership with non-metazoan orthologs reduced the proportion of HGT_C_ to 9.1%, 7.6%, 6.2% and 6.5% for the four bdelloid species, compared with < 1% for all other species (**S5 Data**). The final test was not applicable for the majority of HGT_C_ as orthologs from metazoans were often not detected; thus, the number of genes that additionally showed evidence for monophyly with non-metazoan orthologs was 189, 190, 111 and 82 for *A. ricciae*, *A. vaga*, *R. macrura* and *R. magnacalcarata*, respectively, and was reduced to a handful or zero in all other species (**S5 Data**).

These comparisons support previous findings of a high proportion of non-metazoan genes in bdelloid genomes [26, 28, 40, 93]. Compared to other metazoans, and at all levels of scrutiny, the four bdelloid genomes analysed here showed a substantially greater proportion of genes from non-metazoan sources than do any other species in our comparison. Our results confirm a substantial proportion of foreign genes in the non-desiccating *Rotaria* genomes, in agreement with recent findings based on transcriptomes [40]. Our assessment also showed very low levels of HGT (~1%) into the genome of the anhydrobiotic tardigrade *R. varieornatus,* in agreement with recent estimates [34, 78]. In addition, recent genome investigations of the anhydrobiotic chironomid insect *P. vanderplanki,* which experiences a large number of DNA breakages during desiccation [35], also did not reveal an elevated rate of HGT [94]. Taken together, these findings bring into question the association between anhydrobiosis and elevated rates of HGT that has been previously suggested for bdelloids [26, 28, 30, 31, 93]. One explanation may be that differences in the mechanism of anhydrobiosis, in combination with particular ecological properties of each species, may explain the observed differences in HGT content. Further comparative work is thus required to elucidate any relationship between anhydrobiosis and horizontal transfer.

Alternatively, it is possible that HGT content in bdelloids is impacted as much by a longstanding lack of meiotic sex as by anhydrobiosis. Based on transcriptome data from *Rotaria* species, Eyres *et al.* (2015) have estimated the rate of foreign import to be low in absolute terms, on the order of 13 gains per lineage per million years [40]. This rate may reflect the background rate of initial integration of foreign genes for an organism with similar ecological and physiological properties to bdelloid rotifers. If so, the high proportion of non-metazoan genes encoded in bdelloid genomes may reflect a deviation in the rate of retention, rather than the rate of import. Foreign genes that land in an ameiotic background may persist for longer timescales, even if initially deleterious, given the lack of sexual mechanisms such as segregation that would otherwise rapidly remove them. Thus, foreign genes incorporated by asexuals may tend to accumulate over the extended timescales necessary for domestication.

We also quantified the abundance of transposable elements (TEs) and low-complexity repeats in each genome using a similar approach to that used for the bdelloid genomes (see previous sections and Methods). We chose to focus on the quantification of known repeats and thus did not perform *ab initio* repeat modelling (e.g., RepeatModeler) for non-bdelloid species. There was considerable variation in TE abundance among species (Fig 6E), with the total proportion of genome covered by interspersed repeats varying from 0.3% in the tardigrade *R. varieornatus* to 27.5% in the oyster *Crassostrea gigas* (**S2 Data**). The relative abundance of different classes of repeats, including SINES, LINES, LTR elements and DNA elements also differed greatly among species, as did the amount of simple and low-complexity repeats. The proportion of total repeats (TEs + low-complexity) ranged from 0.6% in *R. varieornatus* to 42% in the annelid worm *Helobdella robusta.* All four bdelloid species display a low abundance of TEs, in agreement with previous findings [23–25, 28, 95]. However, two other species also show a low levels of TEs: *I. linei,* an intracellular parasite of marine invertebrates with a highly reduced genome (42 Mb) [84], and *R. varieornatus,* also with a relatively small genome (56 Mb) [34]. In fact, *R. varieornatus* encodes the fewest TEs of the species analysed here (0.6% as a proportion of assembly span), followed by the four bdelloids (2–3%). These estimates of TE abundances in *I. linei* and *R. varieornatus* are substantially lower than the total repeat content of these genomes (28% and 20% respectively), which includes a high proportion of *ab initio* repeats (inferred directly from the assembled nucleotides) marked as “unclassified” (accounting for ~18% and ~I9% total repeats, respectively [78, 84]), matching our finding of higher *ab initio* repeat content in bdelloids. Additional work is required to elucidate the nature of these unclassified repeats in bdelloids and other taxa.

What evolutionary forces may explain the low abundance of TEs in these species? Asexuality and anhydrobiosis have both previously been posited as factors contributing to the low volume of TEs in bdelloid rotifers. For example, under long-term asexual evolution, TEs may either proliferate freely within a genome and thus drive that lineage to extinction (an extension of Muller’s ratchet), or become lost, domesticated or otherwise silenced [30, 96–98]. Frequent cycles of desiccation and rehydration may also favour the evolution of reduced repeat content, via selection against deleterious chromosomal rearrangements brought about by ectopic recombination of TEs during the repair of DSBs [29, 30].

Our comparisons did not detect any substantial variation in the abundance of known TEs between anhydrobiotic (1.2% and 0.8% for *A. ricciae* and *A. vaga* respectively) versus non-desiccating bdelloids (0.9% and 1.2% for *R. macrura* and *R. magnacalcarata* respectively), despite a considerable increase in the inferred genome size of *Rotaria* species. Moreover, the proposed mechanism involving desiccation relies on DSB repair during rehydration, a process which is presumably limited in the aquatic species *R. macrura* and *R. magnacalcarata* and may also not apply in the case of *R. varieornatus* [34]. However, the majority of bdelloid rotifers are resistant to desiccation, suggesting that anhydrobiosis was probably the ancestral state [17]. Thus, it may be that TEs and other repeats were already largely eradicated in the most recent common ancestor to non-desiccating *Rotaria* species, prior to their adaptation to a fully aquatic lifestyle and loss of anhydrobiosis.

### Concluding remarks

The bdelloid rotifers have drawn attention because two features of their life history are remarkable among metazoans: their apparent ancient asexuality and their ability to withstand desiccation at any life stage. We assessed the relative contributions of asexuality and anhydrobiosis to genome evolution in bdelloids, using de novo whole genome sequence data for desiccating and non-desiccating species and comparisons to both the existing genome of *A. vaga* and to other animal taxa.

We find that both desiccating and non-desiccating species are tetraploid, in agreement with previous work [24, 25], but that homologous divergence in non-desiccating *Rotaria* species is substantially lower than that observed between anhydrobiotic *Adineta* species, and is low compared to estimates of allelic heterozygosity from sexual eukaryotes. This finding runs counter to predictions based on current hypotheses regarding the genomic effects of desiccation, and thus requires a reevaluation of the causes and consequences of intragenomic interactions between bdelloid homologs.

We re-confirm previous reports that bdelloids encode a high proportion of non-metazoan genes. Here too, a role for desiccation-tolerance had been hypothesised. We find that high HGT content is a potentially unique feature of bdelloid genomes among animals, but comparisons to other desiccation-tolerant taxa bring into question the role of anhydrobiosis. Our extensive assembly results also allow for a refinement of the global parameters of bdelloid genomes, and suggest substantial genome size differences between genera that may be linked to desiccation tolerance. The phylogenetic non-independence of these results precludes more direct tests of cause and effect, however, and further genomic evidence is required, ideally from anhydrobiotic species within *Rotaria*.

Comparisons of genome architecture within *Adineta* revealed that a number of unusual genome features reported in *A. vaga,* posited as evidence of long-term ameiotic evolution in this species [28], were largely absent from the closely related species *A. ricciae*, for which a comparable assembly is now available. In addition, we find that bdelloids encode the majority of genes that are required for meiosis and syngamy in sexual taxa, but emphasise that the precise function of these genes in bdelloids is currently unknown. Overall, our results do not exclude the hypothesis that many features of the bdelloid genomes analysed here are consistent with those found in sexual taxa, except for the remarkably high prevalence of HGT.

## Methods

### Rotifer culture and sampling

*Adineta ricciae* [99] rotifers were cultured as previously described [27, 100–102]. Briefly, rotifers were grown in T75 tissue culture flasks (Nunc) with 15–25 ml ddH_2_O, and fed twice a week with 10 μl of either bacteria (*E. coli* TOP10 (ThermoFisher) in water) or a solution of yeast extract and peptone (2.5% w/v each). Approximately 50,000 rotifers were starved overnight before collection and harvested by centrifuging at 10,000 g for 5 mins and discarding the supernatant before treatment according to a DNA or RNA extraction protocol. A starter culture for *R. macrura* was generated from ~100 wild-caught animals isolated from a small pond near Lake Orta, Italy. Populations were grown in sterile distilled water and fed with autoclaved and filter-sterilised organic lettuce extract. Prior to DNA extraction, animals were washed twice in sterile distilled water and starved overnight (~16h), before being washed again with HyPure molecular-grade water. Genomic DNA from approximately 420 animals (260 derived from a single founding animal; the remainder derived from a subpopulation of ~10 wild-caught founders) was extracted using the DNeasy Blood & Tissue kit (Qiagen) following the standard protocol. DNA was extracted in batches and pooled to generate sufficient material. Paired end data for *R. magnacalcarata* have been described previously [40]. Both *R. macrura* and *R. magnacalcarata* PE libraries are derived from multiple-individual samples. For mate-pair library construction for both *R. macrura* and *R. magnacalcarata*, DNA was extracted from a single individual and subjected to whole-genome amplification (WGA) using the Repli-G Single Cell kit (Qiagen), following the manufacturer’s protocol. DNA concentration and quality was ascertained using a Qubit (Invitrogen) and a NanoDrop spectrophotometer (Thermo Scientific).

Desiccation tolerance of *R. macrura* and *R. magnacalcarata* were tested using protocols as previously described [17, 40] (see **S1 Text** for further details).

### Sequencing

For *A. ricciae*, a short-insert library with an insert size of 250 bp was prepared using Illumina Nextera reagents and sequenced (100 bases paired end) on an Illumina HiSeq 2000 at the Eastern Sequence and Informatics Hub (Cambridge, UK). Two long-insert (mate-pair) libraries both with inserts of 3 kb were also sequenced (51 bases paired end) at GATC Biotech (London, UK). In addition, a PacBio (Pacific Biosciences) long read library with an insert of 10 kb was sequenced using 3 SMRT Cells on a PacBio RS II (TGAC The Genome Analysis Centre, Norwich Research Park). An RNA-Seq library with an insert size of 250 bp was sequenced (150 bases paired end) on an Illumina NextSeq500 at the Department of Biochemistry, University of Cambridge (Cambridge, UK). A short-insert library (500 bp insert) for *R. macrura* was prepared using Illumina TruSeq reagents at the Centre for Genomic Research (CGR) at the University of Liverpool (Liverpool, UK). Mate-pair libraries with 2 kb inserts were also prepared at CGR using Nextera reagents, and all libraries were sequenced (I50 bases paired end) over three lanes of an Illumina HiSeq4000 at CGR. Short-insert data for *R. magnacalcarata* has been described previously [40]. All raw data have been submitted to the Sequence Read Archive (SRA), an International Nucleotide Sequence Database Collaboration (INSDC), under the accession IDs ERR2135445–55^2^ (**S1 Table**).

### Genome assembly

For *A. ricciae, R. macrura* and *R. magnacalcarata* data, adapter sequences and low-quality bases were removed from Illumina data using Skewer v0.2.2 [103], and data quality was manually assessed using FastQC v0.11.5 [104]. Genome coverage was estimated by generating kmer distributions using BBMap “kmercountexact” v36.02 [105], and library insert sizes, along with initial genome size estimates, were calculated using SGA “preqc” [106]. Error correction of reads was performed using BBMap “tadpole” (*k* = 31), discarding any pairs of reads containing unique kmers.

Contaminant reads derived from non-target organisms were filtered using BlobTools v0.9.I9 [107]. Briefly, trimmed and error-corrected paired-end data were digitally normalised to ~100X using BBMap “bbnorm” [105] and a preliminary, draft assembly was generated using Velvet v1.2.10 [108], setting a kmer length of 75. Taxonomic annotations for all contigs were determined by comparing contigs against the NCBI nucleotide database (nt) and a custom database containing recently published whole genome sequences of metazoans within the Spiralia (Lophotrochzoa) group (**S4 Table**) using BLAST “megablast” (*E*-value ≤ 1e-25) [109], and the UniRef90 database using Diamond “blastx” [110]. Finally, read coverage for each contig was estimated by mapping non-normalised reads to each draft assembly using BWA “mem” v0.7.12 [111]. Taxon-annotated GC-Coverage plots (“blobplots”) [107,112] were generated using BlobTools (default parameters) and inspected manually. Putative contaminant sequences were identified as contigs showing atypical %GC, read coverage, and/or taxonomic classification. Given the a priori expectation that a substantial number of bdelloid genes may derive from non-metazoan sources, we did not exclude any contigs based on taxonomy alone. Paired reads were excluded from further analysis only if both mapped to an identified contaminant contig, or if one of the pair mapped to a contaminant while the other was unmapped. Additional rounds of filtering were performed if previously unassembled contaminant sequences became evident upon reassembly.

Filtered reads were assembled into contigs using the Platanus assembler v1.2.4 [41] with default parameters. Mate-pair libraries were filtered to remove contaminating FR orientated reads (i.e., reads originating from short fragments) by only including mates where both reads mapped to a distance ≤ 500 bases from the terminus of a contig. Contigs were scaffolded using SSPACE v3.0 [113] and undetermined bases were filled using GapFiller v1.10 [114]. The *A. ricciae* assembly was further scaffolded with the PacBio library using SSPACE-LongRead v1.1 [115]. RNA-Seq reads for *A. ricciae* were assembled de novo using Trinity v2.2.0 [116] (default parameters) and used for additional scaffolding with L_RNA_Scaffolder [117] and SCUBAT v2 [118]. An available transcriptome for *R. magnacalcarata* [40] was similarly utilised. A final round of assembly “polishing” was performed using Redundans v0.12b [42] and scaffolds less than 200 bases in length were discarded. Assembly completeness was evaluated using the Core Eukaryotic Gene Mapping Approach (CEGMA) v2.5 [43] and Benchmarking Universal Single-Copy Orthologs (BUSCO) v3.0.0 [44] gene sets, choosing the Eukaryota (*n* = 303) and Metazoa (*n* = 978) databases in the latter case, and increasing the search limit to eight. Alternative assemblies were also generated using Velvet [108], SPAdes [119] and dipSPAdes [120] for comparison.

The reference assembly pipeline above was designed to maximise assembly contiguity, but may lead to assembly collapse the extent of which is undetermined a priori. Thus, maximum haplotype assemblies were also generated for each species for comparison, defined as assemblies with minimal reduction due to assembly collapse. Maximum haplotype assemblies were generated using either dipSPAdes (default settings) or Platanus with the “bubble crush” reduction parameter set to zero (-u 0). Details of assembly parameters trialled are given in **S1 Data**. Collapsed and maximum haplotype (re)assemblies for *A. vaga* were also generated following the same procedures, using Illumina short-insert libraries (accession IDs SRR801084 and ERR321929) for contig building, and mate-pair (accession ID ERR321928) and 454 data for scaffolding (see [28] for details).

### Gene prediction

Repetitive regions were masked prior to gene prediction. Repeats were modelled *ab initio* using RepeatModeler v1.0.5 [121]. Repeats arising from duplicated genes or recent gene-family expansions (e.g., alpha-tubulin in *R. magnacalcarata* [122]) were removed by comparing each repeat library to the SwissProt database (BLASTX, *E*-value ≤ 1e-5) and retaining only those sequences with descriptions for known repeat elements. The filtered RepeatModeler library was merged with known Rotifera repeats from Repbase v22.02 [123] (accessed using the command “queryRepeatDatabase.pl-species ‘rotifera’’’) and compared to each assembly using RepeatMasker v4.0.7 [124]. Low complexity regions and simple repeats were additionally soft-masked.

Gene prediction was then performed using BRAKER v1.9 [46] where RNA-Seq data was available (*A. ricciae* and *R. magnacalcarata*). Briefly, RNA-Seq reads were aligned to the masked assembly using STAR, specifying the “twoPassMode Basic” parameter to improve splice junction annotation. The resultant alignment BAM file was then input to the BRAKER pipeline with default settings. For *R. macrura*, an initial set of gene models was constructed using MAKER v3.00 [47], using evidence from SNAP [125] and GeneMark-ES v4.3 [126]. MAKER-derived gene models were then passed to Augustus v3.2.1 [48] for final refinement. Transfer and ribosomal RNA genes were predicted using tRNAscan-SE v1.3.1 [127] and RNAmmer [128], respectively. The *A. vaga* 2013 assembly (GCA_000513175.1) was also re-annotated for consistency with these results, using both approaches outlined above (in conjunction with RNASeq library accession ERR260376).

To test if coding sequences had been inadvertently missed during gene prediction, we compared proteins to the source nucleotide sequences from which they had been predicted using TBLASTN (*E*-value ≤ 1e–20). Matches to existing gene models were discounted by removing alignments that showed any overlap with gene regions (BEDtools “intersect” [129] with the “-v” option), leaving only hits to regions of the genome that had not already been annotated as a gene.

### Collinearity analyses

Syntenic regions within and between genomes were identified using MCScanX [49], calling collinear “blocks” as regions with at least five homologous genes and fewer than 10 “gaps” (i.e., missing genes). Rates of synonymous (*K*_s_) and nonsynonymous (*K*_A_) substitution between pairs of collinear genes were estimated by aligning proteins with Clustal Omega [130] and back-translating to nucleotides before calculating *K*_A_ and *K*_s_ values using BioPerl [131]. The collinearity of each block was calculated by dividing the number of collinear genes in a block by the total number of genes in the same region [28]. We also counted the number of collinearity breakpoints between adjacent homologous blocks across each genome, defining a breakpoint as an occurrence where homologous blocks cannot be aligned without rearrangement. All scripts are available at https://github.com/reubwn/collinearity.

### Orthologous clustering and SNP finding

Orthologous relationships among proteins from the same set of protostomes as above were inferred using OrthoFinder v1.1.4 [132] with default settings. All genomic, GFF and protein sequence datasets were downloaded from NCBI Genbank no later than May 2017. For SNP finding, data were mapped using Bowtie2 v2.2.6 [54] with the “--very-sensitive” preset to minimise mismapped reads, and SNPs and indels were called using Platypus v0.8.1 [55], setting a minimum mapping quality of 30, a minimum base quality of 20, filter duplicates = 1 and a minimum read depth to approximately 25% the average coverage of each individual library. VCF manipulation and SNP statistics were calculated using VCFlib vI.0.0-rcI [133]. For *A. vaga*, SNPs were called based on the Illumina dataset ERR321927 mapped to the published genome sequence [28]. For *R. macrura* and *R. magnacalcarata*, SNP_s_ were called based on mate-pair libraries mapped as single-end, as paired-end data for these samples were comprised of multiple non-clonal lineages.

### Horizontal gene transfer

We assessed the extent of horizontal transfer into bdelloid genomes using a combination of sequence comparison and phylogenetics-based approaches, and applied the same tests to a set of 13 publicly available proteomes from species across the Protostomia (**S4 Table**). Protein sequences were first compared to the UniRef90 database [134] (downloaded November 29, 2016) using Diamond “blastp” (*E*-value ≤ 1e-5, maximum target seqs = 500). For each query, two HGT metrics were then calculated: (a) HGT Index (*h_U_* [27]), defined as: *B*_OUT_ – *B*_IN_, where *B*_IN_ is the best (highest) Diamond bitscore from comparisons to “ingroup” taxa and Bout is the corresponding score for hits to “outgroup” taxa; (b) Consensus Hit Support (CHS), defined as the proportion of all hits that support a given query's ingroup/outgroup classification, itself inferred from the highest sum of bitscores to ingroup or outgroup across all hits [92]. The CHS score therefore takes into account the taxonomic distribution of all hits for each query, and militates against misclassifications based on *h_U_* scores alone. We defined the ingroup as “Metazoa” and the outgroup as “non-Metazoa”, and marked all proteins with an *h_U_* ≥ 30 and CHS_OUT_ ≥ 90% as putative HGT_C_ andidates (HGT_C_). We then looked at the distribution of all HGT_C_ across the genome, and discarded any candidate found on a scaffold encoding ≥ 95% genes of putative foreign origin (i.e., “HGT-heavy” scaffolds likely derived from contaminant sequences that were not removed during assembly). For each HGT_C_, physical linkage (i.e., presence on the same scaffold) to a gene with good evidence for metazoan origin (*h*_U_ ≤ 0, CHSin ≥ 90%) and the number of predicted introns was also recorded. Finally, phylogenetic support for HGT was then assessed: for each HGT_C_, the sequences of 15 metazoan and 15 non-metazoan UniRef90 hits (when present) were extracted and aligned using MAFFT v7.309 [135] with default parameters, and a maximum likelihood phylogeny was constructed using IQ-TREE v1.5.3 [136], specifying automatic model selection and 1,000 ultrafast bootstrap replicates. The functionality of GNU Parallel [137] was used to compute multiple trees simultaneously, and clusters with fewer than four taxa were not analysed. Branching patterns of resultant trees were then assessed using a custom script written in R v3.3.1 [138], utilising functions from the “ape” v4.1 package [139]. All HGT analysis scripts are available at https://github.com/reubwn/hgt.

### Transposable elements

The abundance of known transposable elements (TEs) was assessed for the same set of protostomes using RepeatMasker, except using a Repbase (v22.02) repeat library specific to the Metazoa (i.e., “queryRepeatDatabase.pl-species ‘metazoa’’’). Species-specific custom repeat libraries (e.g., using RepeatModeler) were not generated for this analysis; only known repeats from Repbase were compared. The total span of LINEs/SINEs, LTR elements, DNA elements and simple repeats relative to the assembly span for each species was then computed from the RepeatMasker output tables. We also calculated a genome density metric, defined as the number of protein-coding genes per Mb of haploid genome, i.e., accounting for variation in ploidy among species.

### Meiosis genes

The presence of meiosis- and other sex-related genes was assessed following the approach of Tekle *et al*. (2017) [75]. A total of 41 orthologous groups were downloaded from the OrthoMCL database (v5) (http://orthomcl.org/orthomcl/, accessed September 2017) (**S6 Data**). Searches were conducted using both TBLASTN (*E*-value ≤ 1e-5) against the reference assemblies or HMMER3 (http://hmmer.org/) against the predicted protein sets, after alignment with Clustal Omega [130]. Presence was recorded if any query within each OG showed a TBLASTN alignment with ≥ 50% identity over ≥ 50% query length, and/or if HMMER reported an alignment above the significance threshold. The genomes and proteomes of *D. melanogaster* and *C. elegans* were also searched for comparison (**S6 Data**). Proteins were also assessed for presence of the *Zona pellucida*-like domain (Pfam accession PF00100) using HMMER3 as above.

## Acknowledgement

The authors wish to thank Philipp Schiffer and Mark Blaxter (plus lab members) for helpful comments and discussions. We also thank Shilo Dickens at the Department of Biochemistry at the University of Cambridge and the Babraham Institute for help with additional Illumina sequencing. PacBio next-generation sequencing and library construction was delivered via the BBSRC National Capability in Genomics (BB/CCG 1720/1) at Earlham Institute (fka The Genome Analysis Centre) by members of the Genomics Pipelines Group.

## Author contributions

**Conceptualization:** Timothy G. Barraclough, Alan Tunnacliffe.

**Data Curation:** Reuben W. Nowell, Pedro Almeida, Timothy G. Barraclough.

**Formal Analysis:** Reuben W. Nowell, Christopher G. Wilson, Timothy G. Barraclough. Funding Acquisition: Timoth G. Barraclough, Alan Tunnacliffe.

**Investigation:** Reuben W. Nowell, Pedro Almeida, Christopher G. Wilson, Thomas P. Smith, Diego Fontaneto, Alistair Crisp, Gos Micklem, Alan Tunnacliffe, Chiara Boschetti, Timothy G. Barraclough.

**Methodology:** Reuben W. Nowell, Pedro Almeida, Christopher G. Wilson, Alan Tunnacliffe, Chiara Boschetti, Timothy G. Barraclough.

**Project Administration:** Timothy G. Barraclough, Alan Tunnacliffe.

**Resources:** Christopher G. Wilson, Diego Fontaneto, Alan Tunnacliffe, Chiara Boschetti, Timothy G. Barraclough.

**Software:** Reuben W. Nowell, Timothy G. Barraclough.

**Supervision:** Timothy G. Barraclough, Alan Tunnacliffe, Christopher G. Wilson, Chiara Boschetti. Validation: Reuben W. Nowell, Timothy G. Barraclough.

**Visualization:** Reuben W. Nowell.

**Writing—Original Draft Preparation:** Reuben W. Nowell, Timothy G. Barraclough. Writing—Review & Editing: Reuben W. Nowell, Pedro Almeida, Christopher G. Wilson, Diego Fontaneto, Alistair Crisp, Gos Micklem, Alan Tunnacliffe, Chiara Boschetti, Timothy G. Barraclough.

## Supporting Information

**S1 Text.** Test of desiccation tolerance for *Rotaria* species.

**S1 Table.** Data counts and accession numbers for sequence data used in this study.

**S2 Table.** MCScanX collinearity metrics within genomes.

**S3 Table.** Homologous and ohnologous *K*_A_ and *K*_s_.

**S4 Table.** Protostome species included in comparative analysis.

**S5 Table.** OrthoFinder clustering metrics.

**S6 Table.** OrthoFinder clustering metrics per species.

**S1 Fig. Kmer spectra for raw and filtered sequence data.** Distributions show kmer depths *(k* = 31) in raw (gray bars) and filtered (coloured lines) sequence data. Arrows indicate potential secondary and tertiary coverage peaks. The large number of low coverage kmers in the *Rotaria* datasets (particularly *R. magnacalcarata)* indicate substantial levels of polymorphism, most likely due to the sampling approach.

**S2 Fig. Taxon-annotated GC-Coverage plots for initial and final assemblies.** Initial draft assemblies for each species are shown in (i), final reference assemblies in (ii) for (A) *A. ricciae,* (B) *R. macrura* and (C) *R. magnacalcarata*. In each plot, circles represent assembly scaffolds with diameter proportional to the sequence length, positioned by %GC (*X*-axis) and average read coverage (*Y*-axis), and coloured by putative taxon of origin (see legends). Histograms in upper and right panels represent the total span of contigs/scaffolds in a given %GC or coverage bin.

**S3 Fig. Effect of bubble crush on Platanus assembly.**Specifying the removal of polymorphic homologous regions, as identified by Platanus, results in substantial improvements to assembly contiguity by removing a large number of short contigs. Assembly improvements plateau at -u ≥ 0.2. “Bubble span” (third panel) indicates the total length of removed sequences. Values are based on assembly statistics for contigs ≤ 200 bp length.

**S4 Fig. Effect of Redundans reduction.** (a) Identity distribution of discarded sequences to target sequences. (b) Proportion of overlap between discarded and target sequences. (c) Length distribution of discarded sequences.

**S5 Fig. Distribution of intergenic distances between genes.** (A) Box-and-whisker plots showing interquartile range for intergenic distances between genes predicted on the same scaffolds in *A. ricciae* (red), *A. vaga* (orange), *R. macrura* (blue) and *R. magnacalcarata* (green). Median values are indicated by a thick horizontal bar within boxes. Note log_10_ scale on *Y*-axis. (B) Overlayed distributions of intergenic distances for all species. Vertical lines indicate mean distances. Note logm scale on *X*-axis.

**S6 Fig. Distribution of intron length.** (A) Box-and-whisker plots showing interquartile range for all intron lengths in predicted genes from *A. ricciae* (red), *A. vaga* (orange), *R. macrura* (blue) and *R. magnacalcarata* (green). Median values are indicated by a thick horizontal bar within boxes. (B) Distributions of intron size showing that most introns lie in the range 30–100 bp length, but a substantial minority are distributed around a much larger mean (inserts within each panel show distributions for introns > 100 bp length for detail, median values indicated with a black vertical bar). Note log_10_ scale on *X*-axis. The longer average introns in *R. macrura* are driven primarily by an upwards shift in the length of longer introns relative to the other species.

**S7 Fig. MAF distributions for paired-end *Rotaria* sequence data.** A high proportion of low frequency alleles is evident in both mappings, likely caused by population structure in the polyclonal samples (multiple wild-caught individuals) from which these data are derived—the mode at 0.5 is clearer in *R. macrura* as this sample consisted of fewer clones.

**S8 Fig. Coverage profiles for reference and alternative assemblies for *A. vaga*.**
Read coverage distributions for data mapped to the *A. vaga* reference genome GCA_0005 13175.1 (the “2013” assembly) is also bimodal, but in this case the major peak (78.9% sites) is at the lower value centered around ~90x; only a small hump is visible at ~180x. An uncollapsed alternative assembly shows a similar pattern but with the addition of a third coverage peak at approximately 40x (indicated with an arrow). When divergent homologous regions are removed, the majority of sites (65.1 %) show coverage around 180x. Coverage determined from mapping PE library accession ERR321927 (Illumina HiSeq2000, 2 x100bp PE). Values under curves indicate proportion of sites under 50–130x (reference assembly, mode at 90x) and 150–230x (Platanus collapsed assembly, mode at 180x). Only coverage values ≥ 0 are plotted. Mate-pair (accession ERR321928) coverage is also shown.

**S9 Fig. Coverage profiles for reference and alternative *A. ricciae* assemblies.** Paired-end data mapped to both the reference assembly (nAr.v1.8) and alternative assemblies built using Platanus (with polymorphism collapse disabled), Velvet and SPAdes all show similar multimodal coverage distributions, with the majority of sites represented under the higher of the two major coverage peaks, indicating systematic differences in coverage across assemblies regardless of assembly strategy. The coverage profile for the reference genome has modes at approximately 75x (10.5% sites, area under curve from 30 to 90x) and again at 150x (80.8% sites, area under curve from 100 to 200x). Only coverage values ≥ 0 are plotted.

**S10 Fig. Orthologous clustering within bdelloid genomes.** Bars show the distribution of same-species co-orthologous cluster size, expressed as a proportion of each species genome (total number of CDS). Membership size ranges from one (i.e., singletons, left-hand bar) to 10+ (i.e., 10 co-orthologs from the same species, or more; right-hand bar).

**S1 Data.** Assembly metrics for reference and maximum haplotype assemblies.

**S2 Data.** Repeat content for four bdelloid species and 13 other compared protostome species.

**S3 Data.** SNP frequency statistics for detected SNPs across multiple alternative assemblies.

**S4 Data.** Variant Call Format (VCF) files for reads mapped to bdelloid reference assemblies.

**S5 Data.** Horizontal gene transfer metrics for four bdelloid species and 13 other compared protostome species.

**S6 Data.** Meiosis and sex-related gene inventory.

## Footnotes

1 All data are currently under preparation for public submission.

2 All data are currently under preparation for public submission.

